# From goal to outcome: analyzing the progression of biomedical sciences PhD careers in a longitudinal study using an expanded taxonomy

**DOI:** 10.1101/2023.07.13.548881

**Authors:** Abigail M. Brown, Lindsay C. Meyers, Janani Varadarajan, Nicholas J. Ward, Jean-Philippe Cartailler, Roger G. Chalkley, Kathleen L. Gould, Kimberly A. Petrie

## Abstract

Biomedical sciences PhDs pursue a wide range of careers inside and outside academia. However, there is little data regarding how career interests of PhD students relate to the decision to pursue postdoctoral training or to their eventual career outcomes. Here, we present the career goals and career outcomes of 1,452 biomedical sciences PhDs who graduated from Vanderbilt University between 1997-2021. We categorized careers using an expanded three-tiered taxonomy and flags that delineate key career milestones. We also analyzed career goal changes between matriculation and defense, and the reasons why students became more- or less-interested in research-intensive faculty careers. We linked students’ career goal at defense to whether they did a postdoc, the duration of time between defense and the first non-training position, the career area of the first non-training position, and the career area of the job at ten years after graduation. Finally, we followed individual careers for ten years after graduation to characterize movement between different career areas over time. We found that most students changed their career goal during graduate school, declining numbers of alumni pursued postdoctoral training, many alumni entered first non-training positions in a different career area than their goal at defense, and the career area of the first non-training position was a good indicator of the job that alumni held 10 years after graduation. Our findings emphasize that students need a wide range of career development opportunities and career mentoring during graduate school to prepare them for futures in research and research-related professions.

## Introduction

The biomedical research community has been debating the appropriate size and composition of the PhD-trained workforce since Congress passed the National Research Service Awards (NRSA) Act of 1974. The legislation required periodic reviews of the program to ensure it was funding an appropriate number of pre- and postdoctoral trainees to sustain the biomedical research workforce, which was primarily centered in academic institutions at the time ^1^. Indeed, in the early 1980s when the NSF-sponsored Survey of Doctorate Recipients first began tracking tenure-line faculty appointments among U.S. citizens, nearly two-thirds of biomedical PhDs worked in academia and 44% were employed as tenured or tenure-track professors ^2^.

Over the next fifty years, the number of life sciences PhDs working in the U.S. quadrupled without a corresponding increase in employment in faculty jobs ^1, 3^. Much of this growth was absorbed by the private sector, with PhDs finding work in both research- and research-related jobs in the flourishing biotechnology industry. The shifting employment landscape for PhDs, along with a stagnant NIH budget, intensified discussions about the stability and health of the biomedical research workforce and led to a steady drumbeat of calls to action by prominent scientists and research trainees themselves ^4–7^.

Historically, the main source of data about the biomedical research workforce comes from two NSF-sponsored surveys: the Survey of Earned Doctorates given to doctoral students at graduation since 1957, and the longitudinal Survey of Doctoral Recipients, administered since 1973 ^8^. While these surveys provide useful information about the number of PhDs granted and the general types of jobs and sectors in which biomedical PhDs worked, they have numerous limitations ^4, 8^ and do not provide granular data about individual career goals or trajectories. Few institutions have employed systematic approaches to collecting and analyzing data about the careers of their program graduates, and few have adapted their training programs to provide comprehensive career information and preparation to their trainees or meet the needs of the evolving biomedical research workforce.

It is against this backdrop that we began to conduct exit surveys with PhD students at the time of their defense, collecting information about their immediate next steps as well as their long-term career goal. We also began longitudinal tracking of all doctoral alumni who matriculated since 1992 and created a classification system to categorize biomedical PhD career outcomes. Our career outcomes taxonomy resembles recently published classification systems ^9, 10^, with a few notable differences, including the use of novel flags to distinguish key milestones in the career trajectories of alumni (S1 Appendix). We aimed to develop an accurate understanding of how our PhD students’ career plans and careers evolve over time to inform the alignment of our biomedical PhD training programs with workforce needs and to inform our career development initiatives for trainees.

With nearly 25 years of alumni outcomes data and 15 years of PhD student exit survey data in hand, we undertook the present study of the early career trajectories of 1,452 biomedical PhD alumni who matriculated since 1992. We analyzed how career goals changed during graduate school, and the reasons students changed career goals away from, or toward, being an academic PI. We also examined how career goal at the time of defense related to whether alumni did a postdoc or not, and for how long. Consistent with national trends ^11^, we observed a small but steady decline in the percentage of biomedical alumni pursuing postdocs each year. Finally, we followed alumni career trajectories after PhD completion and evaluated how career goal at defense related to their first non-training position and their career area at 1, 3, 5, and 10 years after PhD completion.

## Results

### Career goal at matriculation and defense

Since 2007, we have administered an exit survey to graduates of our institution’s biomedical sciences PhD programs in which they were asked to indicate their primary career goal at two timepoints: the time they matriculated into the PhD program and the time they completed the PhD program (Fig. 1). At matriculation, 62% of students reported being interested in a Research career in the Academic, For-profit, or Government or Nonprofit sectors (n=589). 14% (n=128) of matriculating students indicated they were interested in Teaching careers, and 4% (n=35) were interested in careers categorized as Administrative, Managerial, or Operational jobs (AMO). The remaining students (20%, n=192) were Undecided or Unspecified in their career goal (Und/s).

**Fig 1.**
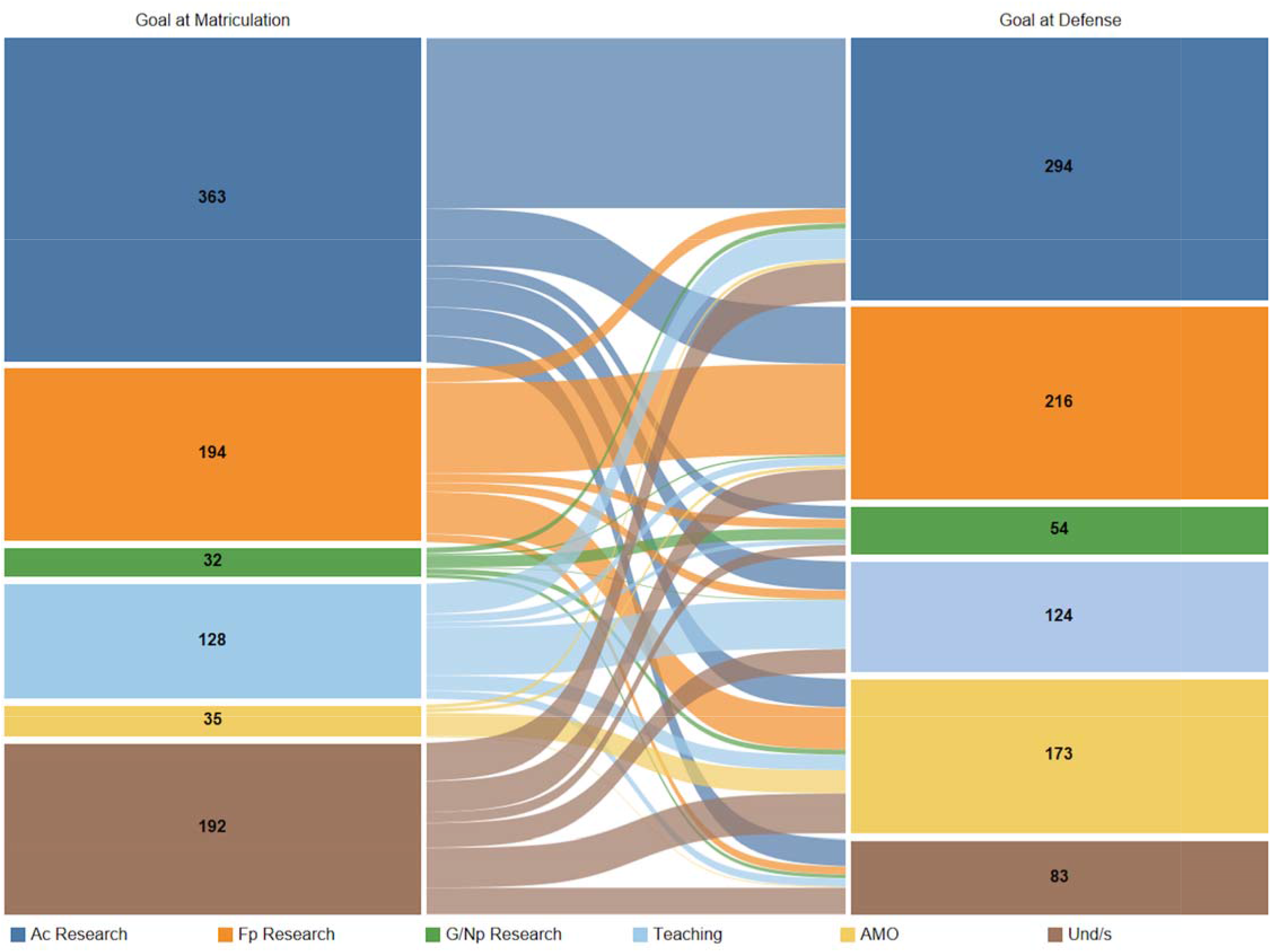
Comparison of biomedical sciences PhD students’ career goal at matriculation and defense. Sankey diagram showing the career goal of biomedical sciences PhD students at matriculation (left nodes) and defense (right nodes). The number of students with a goal in each career area, at matriculation and at defense, is indicated in the corresponding nodes. The links between the nodes show how many students maintained or changed their original career goal. The thickness of each link is proportional to the number of students represented in the link and the colors of the links correspond to the students’ career goal at matriculation. Students who graduated between January 1, 2005, and August 31, 2021, and completed the exit survey were included in this analysis (n=944, Cohort A)

Between matriculation and defense, 56% of all students, including those Und/s at matriculation, had changed their career goal (Fig. 1 and Table 2). 47% of students who initially selected Ac Research changed their primary career goal to another career type, and 153 of these 172 individuals provided free-text comments about why their career goal changed *away* from being a principal investigator (PI) in academia (Table 3). Two-thirds of the reasons cited by students for becoming less interested in PI careers were negative perceptions about academia or being a PI (Table 3: External Factors). Among the most frequently noted reasons were concerns about the competitive funding environment for biomedical research, the lifestyle of academic PIs, and the competition for PI jobs. The remaining one-third of comments were more self-directed factors, such as a perceived mismatch between being a PI and the respondent’s strengths or values or interests, or conversely, a perceived better fit with another type of career. (Table 3: Internal Factors) Other internal sub-themes included exposure to other career options during graduate school, and occasionally, a negative graduate school experience.

**Table 1.**
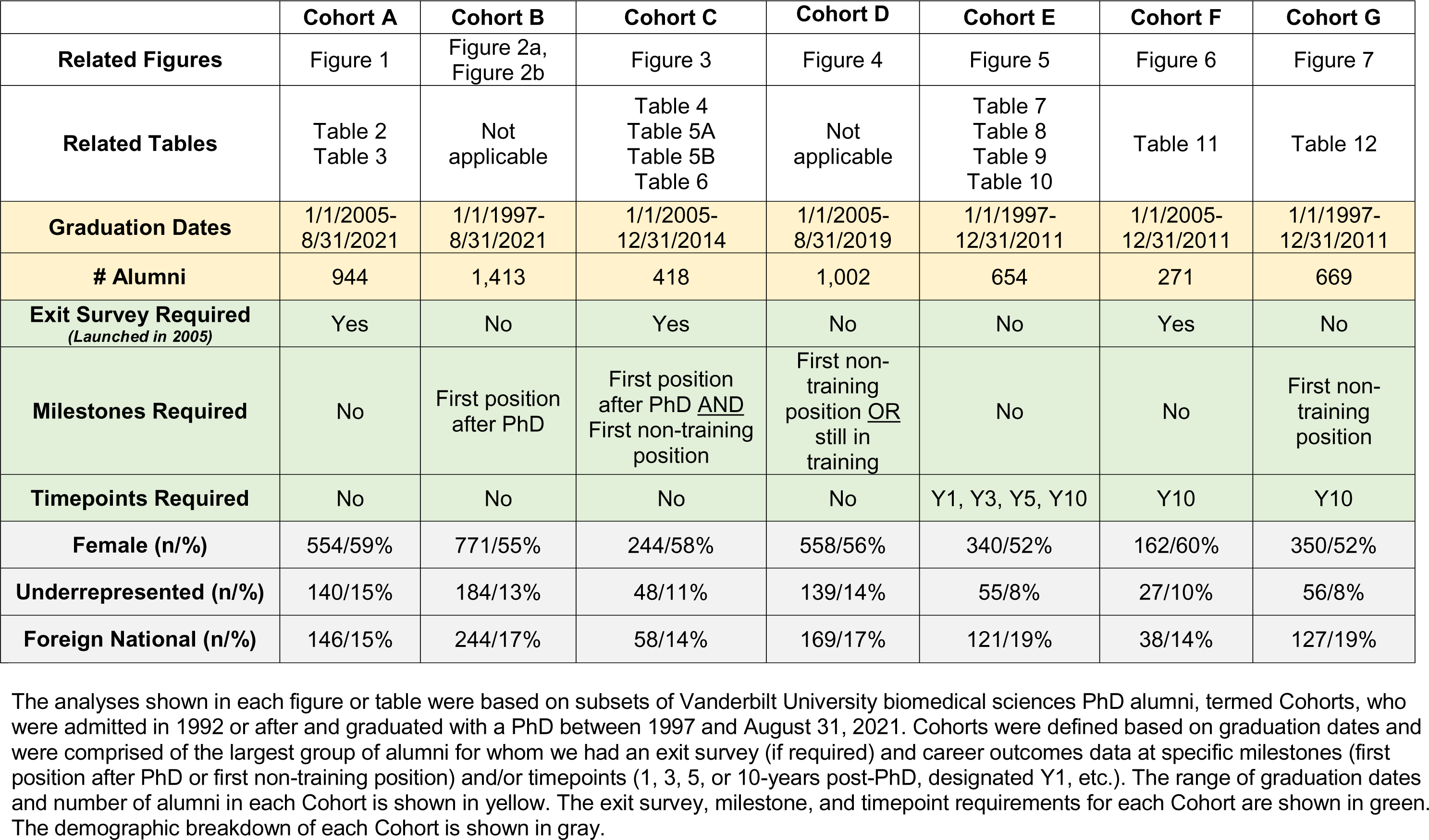
Study cohorts represented in figures and tables.

|  | **Cohort A** | **Cohort B** | **Cohort C** | **Cohort D** | **Cohort E** | **Cohort F** | **Cohort G** |
| --- | --- | --- | --- | --- | --- | --- | --- |
| **Related Figures** | Figure 1 | Figure 2a,<br>Figure 2b | Figure 3 | Figure 4 | Figure 5 | Figure 6 | Figure 7 |
| **Related Tables** | Table 2<br>Table 3 | Not applicable | Table 4<br>Table 5A<br>Table 5B<br>Table 6 | Not applicable | Table 7<br>Table 8<br>Table 9<br>Table 10 | Table 11 | Table 12 |
| **Graduation Dates** | 1/1/2005-8/31/2021 | 1/1/1997-8/31/2021 | 1/1/2005-12/31/2014 | 1/1/2005-8/31/2019 | 1/1/1997-12/31/2011 | 1/1/2005-12/31/2011 | 1/1/1997-12/31/2011 |
| **# Alumni** | 944 | 1,413 | 418 | 1,002 | 654 | 271 | 669 |
| **Exit Survey Required**<br>*(Launched in 2005)* | Yes | No | Yes | No | No | Yes | No |
| **Milestones Required** | No | First position<br>after PhD | First position<br>after PhD <u>AND</u><br>First non-training<br>position | First non-training<br>position <u>OR</u><br>still in<br>training | No | No | First non-training<br>position |
| **Timepoints Required** | No | No | No | No | Y1, Y3, Y5, Y10 | Y10 | Y10 |
| **Female (n/%)** | 554/59% | 771/55% | 244/58% | 558/56% | 340/52% | 162/60% | 350/52% |
| **Underrepresented (n/%)** | 140/15% | 184/13% | 48/11% | 139/14% | 55/8% | 27/10% | 56/8% |
| **Foreign National (n/%)** | 146/15% | 244/17% | 58/14% | 169/17% | 121/19% | 38/14% | 127/19% |

The analyses shown in each figure or table were based on subsets of Vanderbilt University biomedical sciences PhD alumni, termed Cohorts, who were admitted in 1992 or after and graduated with a PhD between 1997 and August 31, 2021. Cohorts were defined based on graduation dates and were comprised of the largest group of alumni for whom we had an exit survey (if required) and career outcomes data at specific milestones (first position after PhD or first non-training position) and/or timepoints (1, 3, 5, or 10-years post-PhD, designated Y1, etc.). The range of graduation dates and number of alumni in each Cohort is shown in yellow. The exit survey, milestone, and timepoint requirements for each Cohort are shown in green. The demographic breakdown of each Cohort is shown in gray.

**Table 2.** PhD students’ career goal at matriculation (rows) and defense (columns).

|  |  | Goal at defense |  |  |  |  |  | Total, goal at matriculation | Percent who changed goal between matriculation and defense |
| --- | --- | --- | --- | --- | --- | --- | --- | --- | --- |
|  |  | Ac Research | Fp Research | G/Np Research | Teaching | AMO | Und/s |  |  |
| Goal at matriculation | Ac Research | 191 | 64 | 14 | 32 | 32 | 30 | 363 | 47% |
|  | Fp Research | 16 | 102 | 10 | 10 | 47 | 9 | 194 | 47% |
|  | G/Np Research | 6 | 2 | 13 | 1 | 6 | 4 | 32 | 59% |
|  | Teaching | 34 | 9 | 5 | 54 | 17 | 9 | 128 | 58% |
|  | AMO | 4 | 4 | - | - | 26 | 1 | 35 | 26% |
|  | Und/s | 43 | 35 | 12 | 27 | 45 | 30 | 192 | 84% |
| Total, goal at defense |  | 294 | 216 | 54 | 124 | 173 | 83 | 944 | 56% of all students |

PhD students who graduated 1/1/2005 to 8/31/2021 and completed the exit survey were included in this analysis (n=944, Cohort A). The shaded number at the intersection of each matching row/column is the number of individuals whose career goal at matriculation matched their career goal at defense. The "Percent who changed goal between matriculation and defense" is the percent of students with a particular goal at matriculation who changed to some other goal by the time of defense, calculated as:

**Table 3.** Frequency with which exit survey respondents cited external or internal themes as reasons for changing their career goal away from or toward becoming a research-focused faculty member in academia.

| **AWAY from being a research-focused faculty member**<br>**(156 comments from 153 individuals)** |  | **TOWARD being a research-focused faculty member**<br>**(88 comments from 68 individuals)** |  |
| --- | --- | --- | --- |
| **External factors** |  | **External factors** |  |
| Lifestyle of PIs | 34 | Positive perceptions of being a PI in academia | 21 |
| Competition for funding | 32 |  |  |
| Competition for faculty jobs | 24 |  |  |
| Academic environment or expectations of PIs | 66 |  |  |
| **Internal factors** |  | **Internal factors** |  |
| Mismatch between academic PI career and skills, values, or interests | 23 | Success in graduate school/enhanced confidence to be successful | 17 |
| Exposure to other career options | 20 | Enjoy research | 14 |
| Match between another career and skills, values, or interests | 15 | Positive mentoring by advisor or others | 13 |
| Negative graduate school experience | 11 | Dislike of other careers or not enough experience to pursue alternatives | 12 |
| **Not descriptive enough to categorize** | 13 | **Not descriptive enough to categorize** | 11 |

PhD students who graduated between 1/1/2005 to 8/31/2021 and completed the exit survey were included in this analysis (n=944, Cohort A). Counts were generated by categorizing free-text responses from individuals who switched their career goal away from, or toward, becoming a research-focused faculty member in academia, and chose to answer the optional question prompt, “If your career goal changed since you started your PhD program, what factor has most affected these goals?” Individual responses were coded across multiple themes if necessary.

There were also 103 students whose primary career goal shifted *toward* becoming an academic PI, 68 of whom provided comments about their reasons. About 25% of reasons involved positive perceptions of the PI role or working in academia such as research independence and lifestyle flexibility (Table 3: External Factors). The remaining 75% of comments related to internal factors; most frequently mentioned were enjoying research, increased confidence about succeeding in an academic career, and encouraging role model/mentor(s).

Similar to Ac Research, 47% of students who were initially interested in Fp Research became interested in another type of career by defense (Table 2). Also, nearly 60% of the students who selected G/Np Research or Teaching at matriculation changed their primary career goal by defense. Persistence of career goal was highest in the AMO group, with only 26% of the 35 students changing their primary goal by defense. 7% of students who entered graduate school interested in a specific career path became Und/s by defense (n=53); most of these students (n=30) switched away from being interested in Ac Research.

As might be predicted, a large number of students who were Und/S about their career goal at matriculation were able to identify a career goal by the time their PhD training ended. Specifically, of the 192 students who were Und/s at matriculation, 84% reported having a defined career goal at defense (Table 2). These students developed an interest in pursuing Ac Research (22%), Fp Research (18%), G/Np Research (6%), Teaching (14%), or AMO (23%).

In summary, a majority of students changed their primary career goal over the course of graduate training. We observed this shift in career goal regardless of the type of career students were interested in at the beginning of graduate school and a similar level of career goal change was observed for most career areas, ranging from 47-59%. Despite this high level of goal switching, the proportion of students interested in some type of Research career remained similar from matriculation (62%) to defense (60%) (Fig. 1). A similar pattern was observed for interest in Teaching careers, where the proportion of students interested in a Teaching career remained nearly identical from matriculation (14%) to defense (13%) even though individual students changed their goal away from, or toward, Teaching. The only career area in which there was a substantial net growth in interest between matriculation and defense was AMO careers, primarily because so few students beginning graduate school were interested in them. Low interest in AMO careers at matriculation is likely due to a lack of familiarity with AMO careers compared to the teaching and research careers that PhD students universally encountered during college and pre-graduate school research experiences.

### First position after PhD

Traditionally, it has been the norm for biomedical PhD graduates to enter a postdoctoral research position immediately after completion of the PhD. Out of 1,413 students who graduated between 1997 and August 31, 2021, 75% entered postdoctoral training as their first position (Fig. 2a). 84% of postdoctoral positions were in Ac Research, 14% were in G/Np Research, 2% were in Fp Research, and three individuals entered postdoctoral positions focused on Teaching or AMO (0.3%). An additional twelve individuals eventually did one or more postdocs, primarily in Ac Research, but it was not the first position they held after completing their PhD.

**Fig 2.**
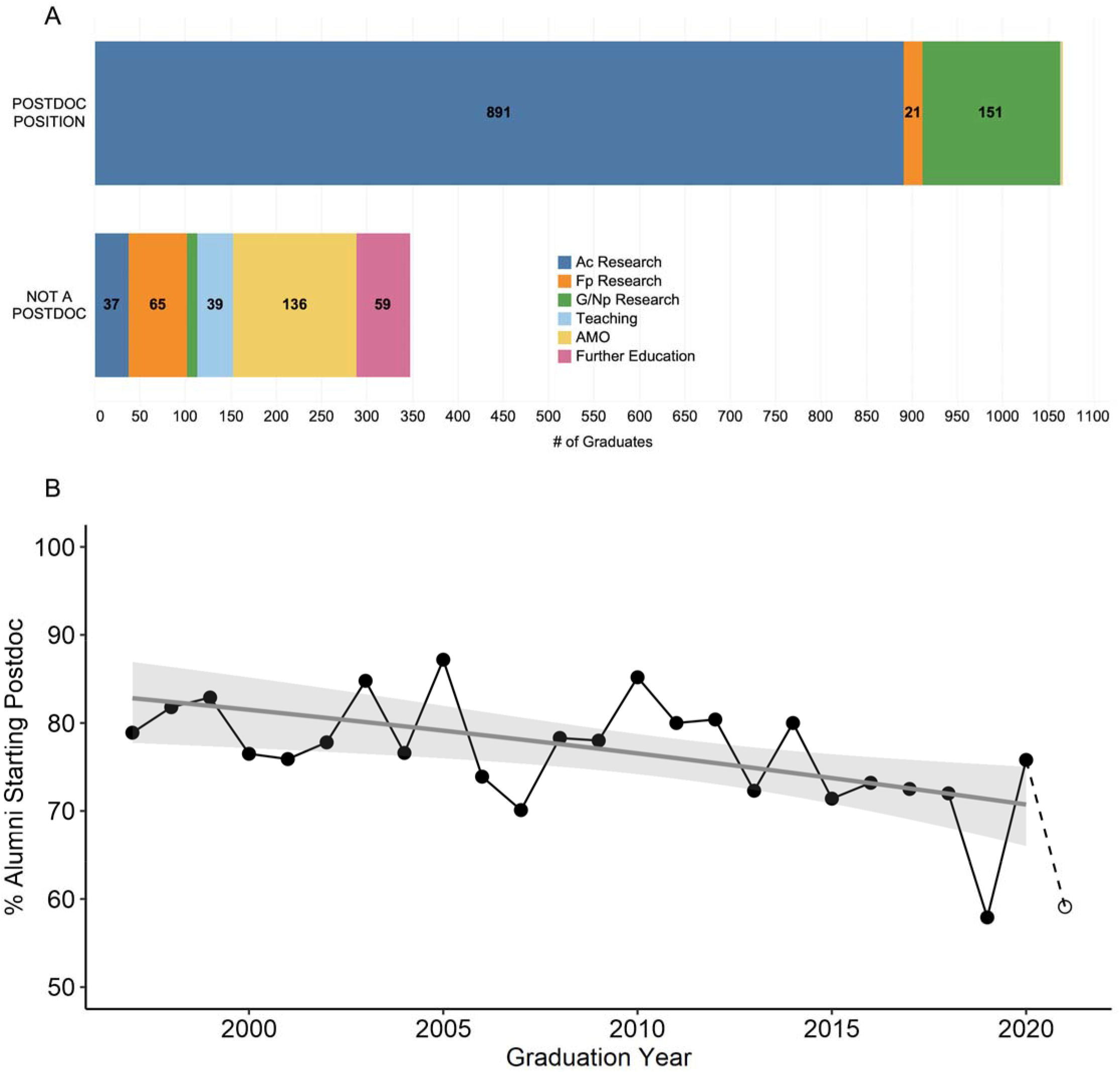
First position after PhD completion. A) Stacked bar chart showing the career area of alumni who did (top) or did not do (bottom) a postdoc as their first position after PhD completion. Students who graduated between 1/1/1997 and 8/31/2021 and completed the exit survey were included in this analysis (n=1,413, cohort B). B) Line graph (black) indicating percent of biomedical sciences PhD alumni who conducted a postdoc immediately after PhD completion, grouped by year of graduation. Full-year data (1997-2020) are represented with filled dots, and half-year data (2021) are represented with open dots. The probability of alumni conducting a postdoc as their first position after PhD completion was modeled using simple logistic regression (gray line with shaded 95% confidence interval). Students who graduated 1/1/1997-8/31/2021 were included in the line graph (n=1,413, cohort B), and students who graduated 1/1/1997-12/31/2020 were used for logistic regression (n=1,369, full-year subset of cohort B).

The next steps of the 347 alumni who did not immediately do a postdoc were varied. 17% of these individuals pursued further educational degrees in law, medicine, nursing, veterinary medicine, or business, and the remainder entered employment directly after graduating. Of the 288 who took jobs, 11% entered Ac Research, 19% entered Fp Research, 3% entered G/Np Research, 11% entered Teaching, and 40% entered AMO positions.

Between 1997-2021, there was year-to-year variability in the percentage of graduates who pursued postdoctoral training as a first position after PhD completion (Fig. 2b). Logistic regression was conducted on full-year data (1997-2020) to create a model estimating the probability that a graduate began postdoctoral training. The model presented here, which incorporated graduation year as a predictor of postdoctoral training, fit these data better than an "intercept-only" model that did not use graduation year as a predictor (χ^2^(1) = 8.22, p = 0.004, likelihood ratio test). The odds ratio corresponding to graduation year was 0.97 with 95% confidence interval (CI) [0.95, 0.99]. The upper limit of this 95% CI is less than 1, indicating the odds of a graduate entering postdoctoral training decreased as graduation year increased. From our model, the probability of entering postdoctoral training started at 83% with 95% CI [78%, 87%] in 1997 and dropped to 71% with 95% CI [66%,75%] by 2020. Based on this modeling, the probability of students entering postdoctoral training decreased by approximately 15% over time from 1997 to 2020.

### Career goal and first non-training position

To characterize the early career paths of our alumni, we related their career goal at defense to their first position after PhD completion and their first non-training position (Fig. 3 and Tables 4, 5A, 5B). We restricted this analysis to alumni who graduated before 2015 to avoid biasing the cohort toward individuals who entered AMO or Teaching for their first non-training position, as these individuals were more likely to do short postdocs or skip postdoctoral training altogether. There were 418 alumni who graduated between 2005-2014 for whom we had all three of these data points. Overall, 78% of this cohort of 418 alumni did a postdoc and 22% did not.

**Fig 3.**
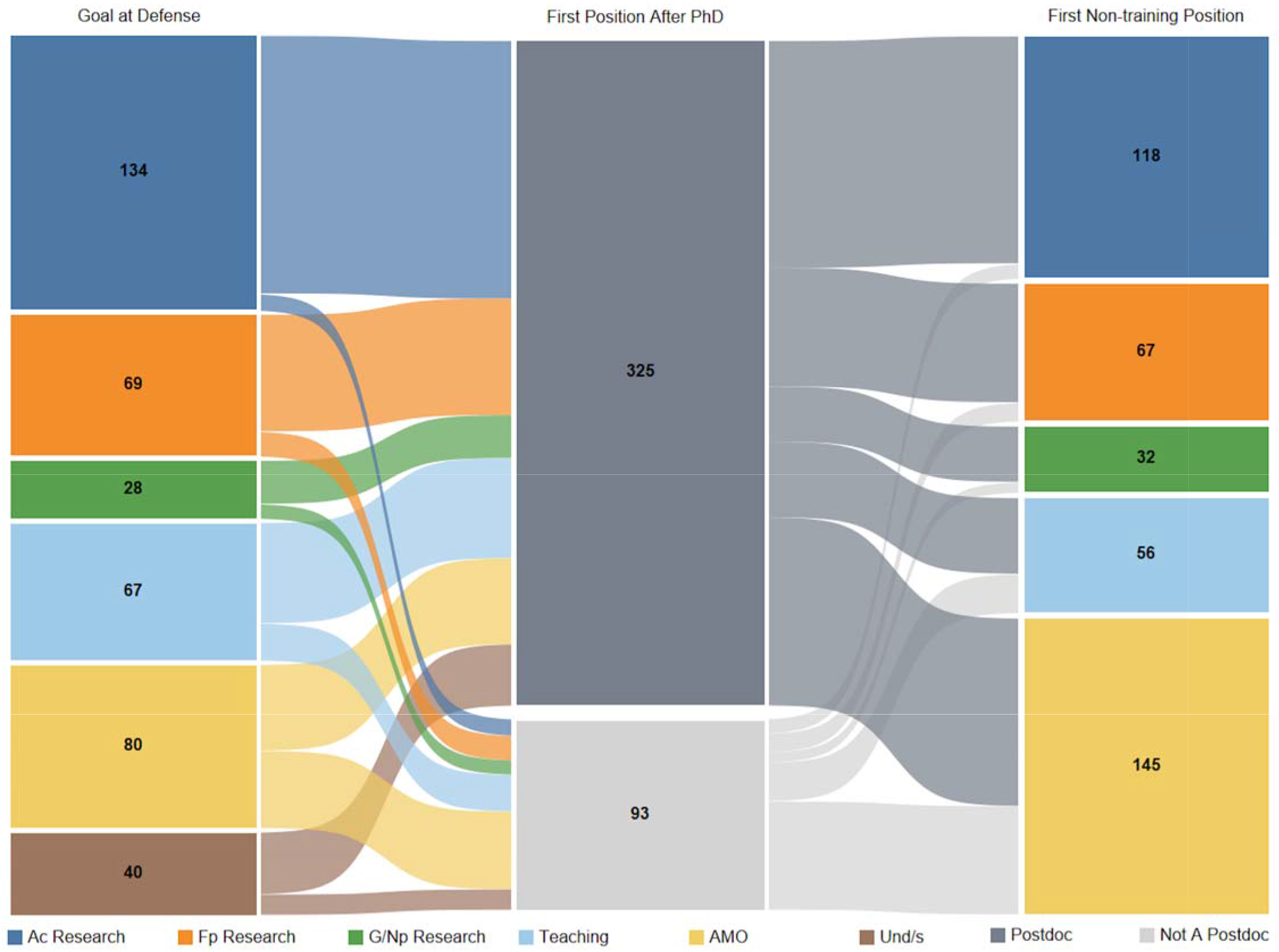
Comparison of career goal at defense with first position after PhD completion and first non-training position. Sankey diagram showing the career goal of biomedical sciences PhD students at defense (left nodes), first position after PhD completion (middle nodes, grayscale), and first non-training position (right nodes). The links between the left and the middle nodes represent the movement of alumni from each career goal area to their first position after PhD completion, categorized as either a postdoc or not a postdoc position. The links from the middle to the right nodes show the movement of alumni from their first position after PhD completion to their first non-training position. The thickness of each link is proportional to the number of students represented in the link and the colors of the links between two nodes correspond to the color of the respective career category of the earlier node. For the 93 individuals who did not do a postdoc (middle node, light gray), their first position after PhD was their first non-training position. Students who graduated from 1/1/2005 to 12/31/2014 and completed the exit survey were included in this analysis (n=418, cohort C).

**Table 4.**
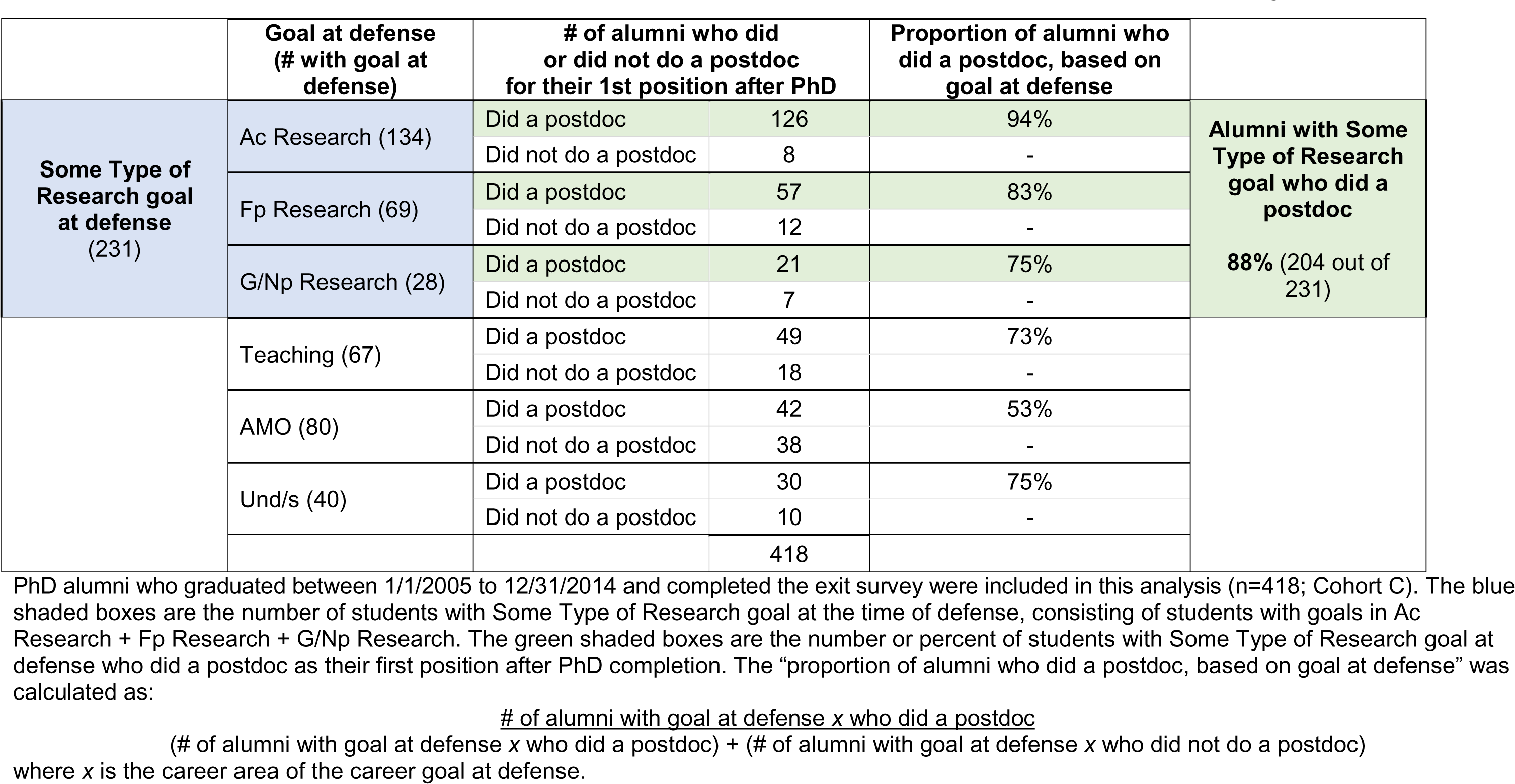
Proportion of PhD alumni who did a postdoc for their first position after PhD completion, based on career goal at defense.

We found that participation in postdoctoral training correlated with career goal at defense. Almost all individuals with some type of Research career goal at defense pursued postdoctoral training (88%; 204 of 231; Fig. 3 and Table 4). Further, as shown in Table 5A, 71% of this cohort who did a postdoc went on to some type of Research job for their first non-training position (144 of 204), while 6% (13 of 204) entered a Teaching position, and 23% entered an AMO job upon completing their postdoctoral training (47 of 204).

**Table 5A.** First non-training position of alumni who did a postdoc as their first position after their PhD, based on their career goal at the time of defense.

|  | **Goal at defense**<br>(# with goal at defense<br>who did a postdoc) | **# of alumni in each career**<br>**area for 1<sup>st</sup> non-training**<br>**position** |  | **Proportion of alumni in career area**<br>**of goal at defense for 1<sup>st</sup> non-**<br>**training position** |
| --- | --- | --- | --- | --- |
| **Some Type of**<br>**Research goal at**<br>**defense, and did**<br>**a postdoc (204)** | Ac Research (126)<br>Fp Research (57)<br>G/Np Research (21) | Some Type of Research<br>*Ac Research (80)*<br>*Fp Research (44)*<br>*G/Np Research (20)* | 144 | 71% |
|  |  | Teaching | 13 | 6% |
|  |  | AMO | 47 | 23% |
|  | Teaching (49) | Some Type of Research<br>*Ac Research (13)*<br>*Fp Research (3)*<br>*G/Np Research (2)* | 18 | 37% |
|  |  | Teaching | 20 | 41% |
|  |  | AMO | 11 | 22% |
|  | AMO (42) | Some Type of Research<br>*Ac Research (6)*<br>*Fp Research (6)*<br>*G/Np Research (2)* | 14 | 33% |
|  |  | Teaching | 2 | 5% |
|  |  | AMO | 26 | 62% |
|  | Und/s (30) | Some Type of Research<br>*Ac Research (12)*<br>*Fp Research (5)*<br>*G/Np Research (3)* | 20 | 67% |
|  |  | Teaching | 2 | 7% |
|  |  | AMO | 8 | 27% |

Career progression of PhD alumni in Cohort C (graduated 1/1/2005 to 12/31/2014 and completed the exit survey; n=418) who did a postdoc as their first position after PhD completion (204 of 418). The blue shaded boxes are the number of students with Some Type of Research goal at defense who did a postdoc, consisting of students with goals in Ac Research + Fp Research + G/Np Research. The green shaded boxes are the number of alumni in Some Type of Research role for their 1<sup>st</sup> non-training position. The “proportion of alumni in career area of goal at defense for 1st non-training position” was calculated as:

73% of individuals interested in Teaching at defense did a postdoc as their first position after PhD completion (49 of 67; Fig. 3 and Table 4). As shown in Table 5A, 41% of these individuals (20 of 49) entered a Teaching job for their first non-training position, but 37% (18 of 49) entered a Research career of some type and 22% (11 of 49) entered an AMO job as their first non-training position. Another group with high participation in postdoctoral training comprised individuals who were Und/s at the time of defense; 75% of this group (30 of 40) also did a postdoc (Table 4), and 67% of these went on to a research career (20 of 30; Table 5A).

Those with AMO career goals at the time of defense were the least likely to pursue postdoctoral training, but 53% of these individuals (42 of 80) still opted to do a postdoc (Table 4). Of these 42 individuals, 62% (26) eventually entered an AMO career, 33% (14) entered some type of Research job, and 5% (2) started a Teaching job after completing postdoctoral training (Table 5A).

For alumni who entered postdoctoral training, the length of time between doctoral thesis defense and the first non-training position varied by students’ career goal at defense (Fig. 4A). This analysis was intentionally limited to those alumni who entered a postdoc immediately after PhD completion, allowing this measure to serve as an approximation of total time spent in postdoctoral training. Alumni with Ac Research or Und/s career goals at defense had the longest time to first non-training position (median length: 5.5 and 4.3 years, respectively), and those with AMO career goals had the shortest (median length: 3.0 years). Alumni who did not take an exit survey at defense had a median time to first non-training position of 4.3 years, likely representing a mixture of multiple career goals (data not shown). Comparing the cumulative incidence curves, individuals with Fp Research, G/Np Research, Teaching, AMO, or Und/s goals at defense tracked together for the first few years after thesis defense. By 2.5 years after doctoral defense, those interested in Ac Research careers exhibited a divergence toward staying in training longer than other groups. Around the three-year timepoint after defense, the curve for those with AMO goals began to diverge toward a greater probability of entering a first non-training position. Shortly after the four-year timepoint, all groups but Ac Research reached their median values (50% probability of starting a first non-training position), indicating individuals interested in Ac Research careers tended to spend a longer time in training than those interested in other career areas. Interestingly, the cumulative incidence curve of those who were Und/s tracked initially with those who had non-Ac Research goals at defense, but the Und/s curve diverged around the two-year timepoint until it became aligned around the six-year timepoint with those who had Ac Research goals at defense, suggesting that some who were Und/S at defense were considering an Ac Research career. Out of all pairwise comparisons, the cumulative incidence curves for Ac Research and AMO exhibited the least overlap in confidence bands across all time durations, with the tail of the Ac Research curve extending into longer training times (Fig. 4B).

**Fig 4.**
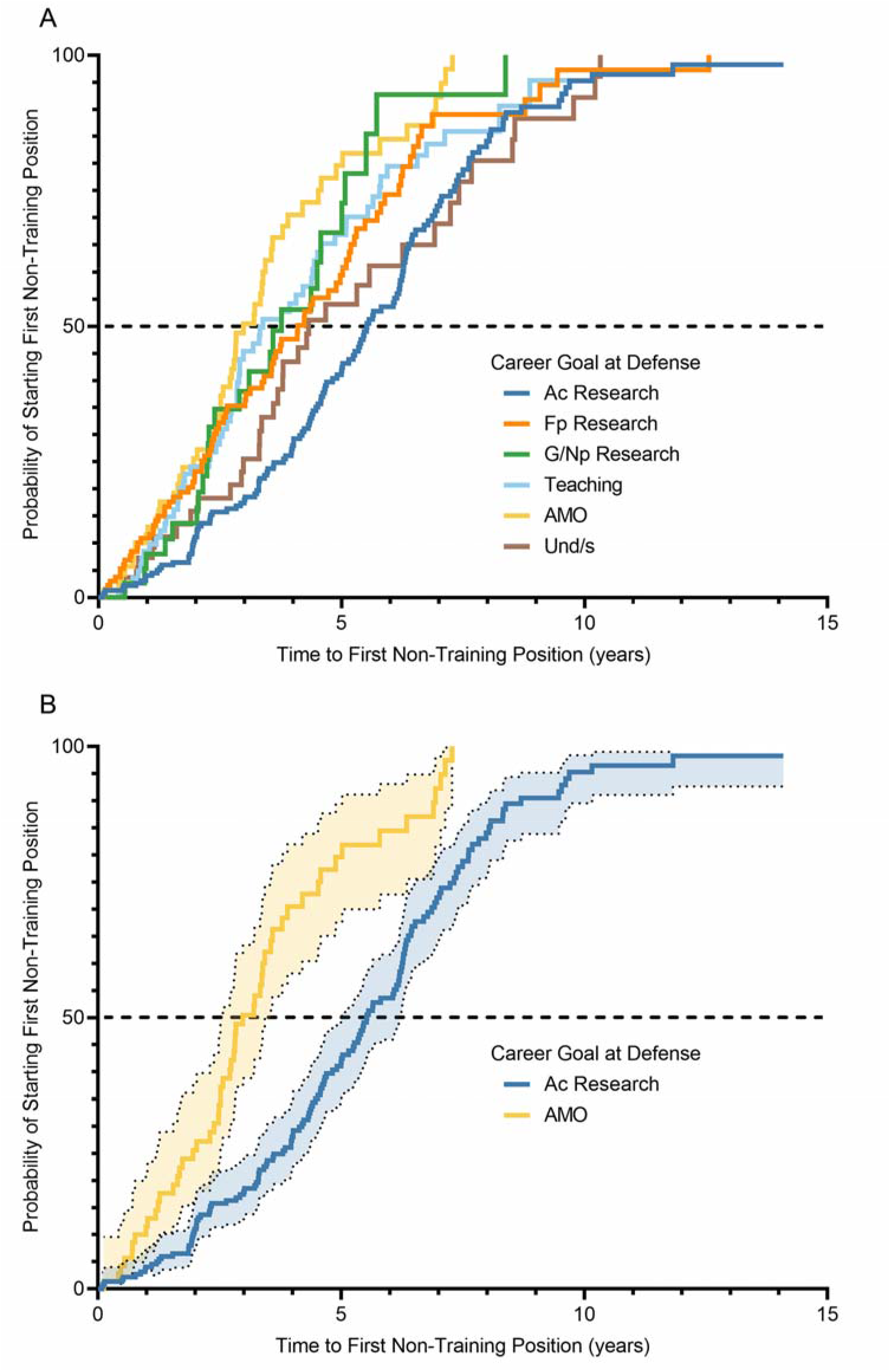
Cumulative incidence curves showing the estimated probability of entering a first non-training position at timepoints following doctoral thesis defense. Curve color corresponds to career area designated in legend with graphs showing A) all career areas, and B) Ac Research vs AMO with 95% confidence bands. Alumni who graduated 1/1/2005-8/31/2019 were included in the probability calculations if they entered a postdoctoral training position immediately after their thesis defense and had either i.) completed training and entered their first non-training position, or ii.) were still in training and had not entered a first non-training position at the cut-off date for recorded observations (a subset of Cohort D, n=773). The 229 individuals in Cohort D who did not do a postdoc immediately after defense but instead entered a first non-training position were not included in the analysis. Individuals per career goal at defense and median time after defense to first non-training position (time at which curve crosses 50% probability threshold): Ac Research (n=230; 5.5 years), Fp Research (n=132; 4.1 years), G/Np Research (n=40; 3.8 years), Teaching (n=84; 3.4 years), AMO (n=71; 3.0 years), Und/s (n=57; 4.3 years).

Ninety-three alumni in this cohort of 418 graduates (cohort C) chose not to do a postdoc (Fig. 3 and Tables 4 and 5B). As shown in Table 5B, 53 went directly into an AMO career area, including 34 of 38 individuals (89%) whose stated career goal at defense was an AMO career. Nineteen alumni started a Teaching position immediately after PhD completion, including 13 of 18 individuals (72%) whose stated career goal was a Teaching career. Interestingly, 21 alumni entered a Research job immediately after PhD completion instead of doing a postdoc: 7 in Ac Research, 9 in Fp Research, and 5 in G/Np Research.

**Table 5B.** First non-training position of alumni who did <u>not</u> do a postdoc as their first position after PhD completion, based on their career goal at the time of defense.

|  | **Goal at defense**<br>(# with goal at graduation who did not do a postdoc) | **# of alumni in each career area for 1<sup>st</sup> non-training position** |  | **Proportion of alumni in career area of goal at defense for 1<sup>st</sup> non-training position** |
| --- | --- | --- | --- | --- |
| **Some Type of Research goal at defense, and did not do a postdoc (27)** | Ac Research (8)<br>FP Research (12)<br>G/Np Research (7) | Some Type of Research<br>*Ac Research (6)*<br>*Fp Research (6)*<br>*G/Np Research (2)* | 14 | 52% |
|  |  | Teaching | 3 | 11% |
|  |  | AMO | 10 | 37% |
|  | Teaching (18) | Some Type of Research<br>*Ac Research (0)*<br>*Fp Research (0)*<br>*G/Np Research (1)* | 1 | 6% |
|  |  | Teaching | 13 | 72% |
|  |  | AMO | 4 | 22% |
|  | AMO (38) | Some Type of Research<br>*Ac Research (0)*<br>*Fp Research (1)*<br>*G/Np Research (2)* | 3 | 8% |
|  |  | Teaching | 1 | 3% |
|  |  | AMO | 34 | 89% |
|  | Und/s (10) | Some Type of Research<br>*Ac Research (1)*<br>*Fp Research (2)*<br>*G/Np Research (0)* | 3 | 30% |
|  |  | Teaching | 2 | 20% |
|  |  | AMO | 5 | 50% |

Career progression of PhD alumni in Cohort C (graduated 1/1/2005 to 12/31/2014 and completed the exit survey; n=418) who did not do a postdoc as their first position after PhD completion (93 of 418). The blue shaded boxes are the number of students with Some Type of Research goal at defense who did not do a postdoc, consisting of students with goals in Ac Research + Fp Research + G/Np Research. The green shaded boxes are the number of alumni in Some Type of Research role for their 1<sup>st</sup> non-training position. The “proportion of alumni in career area of goal at defense for 1st non-training position” was calculated as:

While there is considerable similarity between the relative distribution of students’ career goal at defense and the relative distribution of first non-training positions (Fig. 3), a direct comparison of individuals’ career goal at defense to their first non-training position revealed that only 51% of graduates entered a first non-training position that was the same as their career goal at defense (Table 6). The alignment of career goals at defense and first non-training position was highest among those interested in AMO careers, with 75% of individuals who were interested in an AMO career at defense entering an AMO career as a first non-training job. About half of those interested in either Ac Research or Teaching careers at defense entered those career paths for their first non-training position (52% and 49%, respectively). Only 35% of those interested in Fp Research careers, and only 21% of those interested in G/Np Research careers, entered those types of jobs for their first non-training role. Overall, the match between career goal at defense and first non-training position was lower for those who did a postdoc compared to those who did not (45% vs. 71%, respectively; data not shown). Taken together, the results suggest that goals at defense are not a reliable indicator of the career area of the first non-training position, especially when there is an intervening period of postdoctoral training.

**Table 6.** Goal at defense compared to first non-training position.

|  |  | 1st non-training position |  |  |  |  | Total,<br>goal at defense | % of alumni whose 1<sup>st</sup> non-training position matched<br>their goal at defense |
| --- | --- | --- | --- | --- | --- | --- | --- | --- |
|  |  | Ac<br>Research | Fp<br>Research | G/Np<br>Research | Teaching | AMO |  |  |
| Goal at defense | Ac Research | 69 | 22 | 11 | 9 | 23 | 134 | 51.5% |
|  | Fp Research | 15 | 24 | 5 | 5 | 20 | 69 | 35% |
|  | G/Np Research | 2 | 4 | 6 | 2 | 14 | 28 | 21% |
|  | Teaching | 13 | 3 | 3 | 33 | 15 | 67 | 49% |
|  | AMO | 6 | 7 | 4 | 3 | 60 | 80 | 75% |
|  | Und/s | (13) | (7) | (3) | (4) | (13) | (40) | Not applicable |
| Total, 1st non-training position |  | 105<sup>†</sup> | 60<sup>†</sup> | 29<sup>†</sup> | 52<sup>†</sup> | 132<sup>†</sup> | 378<sup>†</sup> | 51% across all career areas<sup>†</sup> |

PhD students who graduated 1/1/2005 to 12/31/2014 and completed the exit survey were included in this analysis (n=418; Cohort C). The shaded number at the intersection of each matching row/column is the number of alumni whose career goal at defense matched the career area of their first non-training position (192 individuals across all career areas). Alumni with an Und/s career goal at defense are shown in parentheses because they were excluded from the totals prior to the calculation of percentages. <sup>†</sup>indicates totals that excluded individuals with Und/s career goals at defense. The % of alumni whose 1<sup>st</sup> non-training position matched their goal at defense was calculated as:

### Careers during first 10 years after PhD

By following individual careers longitudinally, we were able to see how the careers of 654 alumni progressed throughout the ten years following graduation with a PhD (Fig. 5, cohort E). At one year after graduation (Y1), 64% of alumni were in Ac Research. During the following nine years, 59% of those who initially worked in Ac Research (246 out of 418) and 88% of those who initially worked in G/Np Research (75 out of 85) transitioned to other career types. The gradual attrition from Ac or G/Np Research careers to other career types parallels the movement of alumni out of postdoctoral training into jobs, a transition that is largely complete by five years after graduation (Y5; Table 7). Despite this attrition, 53% of alumni were still employed in some type of Research career at Y10, with the majority of these in Ac Research, and 24% held tenure-track faculty appointments by Y10. About two-thirds of tenure-track positions were research-focused positions in academia, government, or nonprofit organizations and one-third were teaching-focused positions (Table 8).

**Fig 5.**
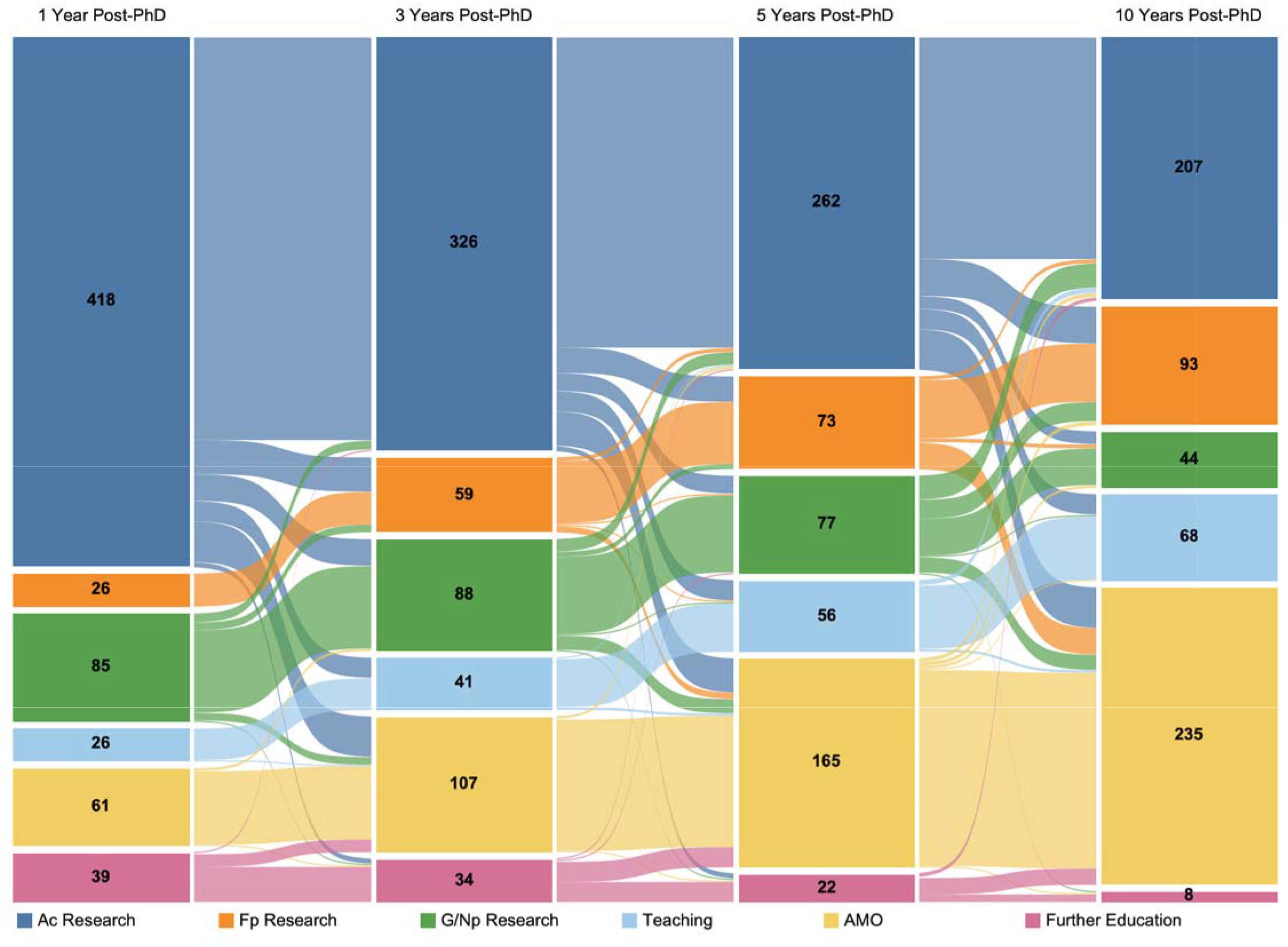
Movement of PhD alumni between different career areas in the first 10 years after graduation. Sankey diagram showing the career area of biomedical sciences PhD alumni at four timepoints from one to ten years after graduation. Nodes show the career area at each timepoint, and links show the movement of alumni between career areas. The color of the link between two timepoints corresponds to the career area at the earlier timepoint and the thickness of each link is proportional to the number of alumni represented in the link. Students who graduated 1/1/1997 to 12/31/2011 were included in this analysis (n=654, cohort E).

**Table 7.**
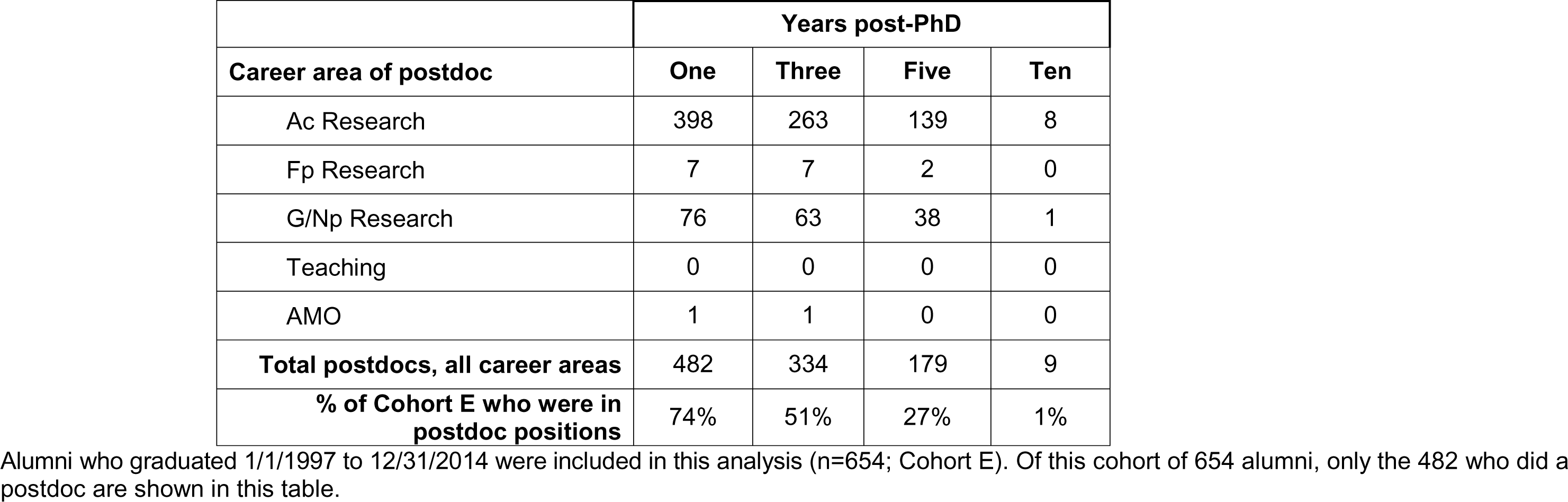
Count of postdocs, by career area of postdoc, at 1, 3, 5, or 10 years post-PhD.

**Table 8.**
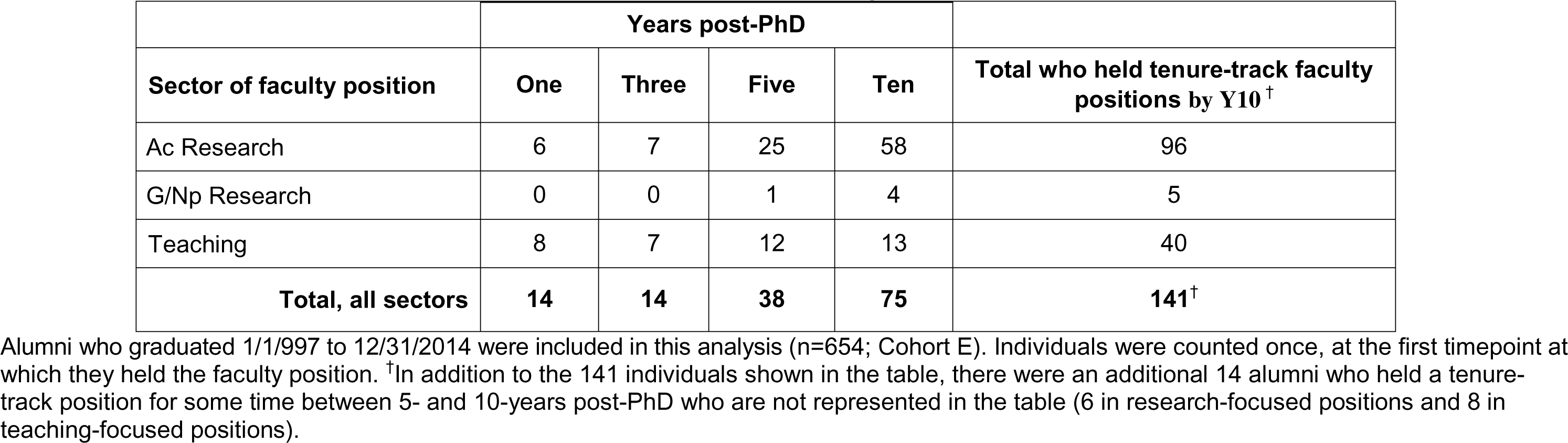
Timepoint at which alumni first held a tenured or tenure-track faculty position.

|  | Years post-PhD |  |  |  |  |
| --- | --- | --- | --- | --- | --- |
| Sector of faculty position | One | Three | Five | Ten | Total who held tenure-track faculty positions by Y10<sup>†</sup> |
| Ac Research | 6 | 7 | 25 | 58 | 96 |
| G/Np Research | 0 | 0 | 1 | 4 | 5 |
| Teaching | 8 | 7 | 12 | 13 | 40 |
| **Total, all sectors** | **14** | **14** | **38** | **75** | **141<sup>†</sup>** |

Alumni who graduated 1/1/1997 to 12/31/2014 were included in this analysis (n=654; Cohort E). Individuals were counted once, at the first timepoint at which they held the faculty position. <sup>†</sup>In addition to the 141 individuals shown in the table, there were an additional 14 alumni who held a tenure-track position for some time between 5- and 10-years post-PhD who are not represented in the table (6 in research-focused positions and 8 in teaching-focused positions).

When alumni were grouped by career area at Y10, individuals who were employed in Ac Research, Fp Research, and G/Np Research were significantly more likely to have spent a longer time in postdoc positions than those who were employed in AMO at Y10 (Table 9 lists p values for significant pairwise comparisons; H(4) = 69.59, p < 0.0001, Kruskal-Wallis test). Likewise, those employed in Ac Research at Y10 were significantly more likely to have spent a longer time in postdoc positions than those employed in Teaching at Y10 (Table 9). Overall, those employed in Ac Research positions at Y10 had the longest median postdoctoral training (4.9 years) and those employed in AMO positions at Y10 had the shortest (2.8 years).

**Table 9.** Duration of postdoctoral training of individuals who conducted postdocs, based on career outcome at Y10.

| **Career area ten years post-PhD** | **# of alumni who pursued postdoctoral training** | **Median length of postdoc, years (interquartile range)** | **Statistically significant comparisons** |
| --- | --- | --- | --- |
| Ac Research | 190 | 4.9 (3.1 – 6.1) | vs. Teaching ($p = 0.0004$)<br>vs. AMO ($p < 0.0001$) |
| Fp Research | 81 | 4.4 (2.8 – 6.2) | vs. AMO ($p < 0.0001$) |
| G/Np Research | 37 | 4.1 (2.4 – 6.0) | vs. AMO ($p = 0.0036$) |
| Teaching | 50 | 3.2 (1.8 – 5.1) | vs. Ac Research ($p=0.0004$) |
| AMO | 151 | 2.8 (1.6 – 4.3) | vs. Ac Research ($p<0.0001$)<br>vs. Fp Research ($p<0.0001$)<br>vs. G/Np Research ($p=0.0036$) |
| **All career areas** | **509** | **4.0 (2.2 – 5.8)** |  |

Alumni who graduated 1/1/1997 to 12/31/2011 were included in this analysis ( $n=654$ ; Cohort E). From this cohort, only those individuals who entered postdoctoral training as the first position after PhD completion were included in the calculation of median length of postdoc based on Y10 career outcome ( $n=509$ of 654). Pairwise comparisons were made between all groups using Dunn's multiple comparisons testing, and the four statistically significant pairwise comparisons observed are noted in the table.

During the ten years following graduation, movement out of, or between, different types of Research careers was more common than movement out of Teaching or AMO careers, where alumni tended to remain once they entered jobs in those areas. There was a net growth in employment in Fp Research careers from 4% to 14% (Fig. 5), but overall alumni tended to migrate from Research careers into AMO careers at each timepoint examined. Since transitions out of Teaching or AMO careers were infrequent compared to transitions into these career areas, employment in Teaching careers grew from 4% to 10%, employment in AMO careers grew from 9% to 36% as alumni gained experience and started to move away from the bench into managerial roles, and the number of alumni pursuing further education decreased over time.

To gain insight into how alumni careers evolve, we examined the proportion of alumni at each timepoint whose career area was identical to the career they held at Y10 after graduation. Across all career areas, a progressively greater proportion of alumni were employed in the same career area at Y1, Y3, and Y5, respectively, as the job they would eventually hold at Y10 (Table 10 and S3 Table). At Y1, 42% of alumni were employed in the same career area in which they would be employed in Y10; this grew to 54% and 69% at Y3 and Y5, respectively. Notably, 80% of alumni who were employed in a Teaching career at Y1, and nearly 100% of alumni who were employed in an AMO career at Y1, were still employed in those career areas at Y10. The proportion of individuals who would remain employed in some type of Research career area at Y10 grew from 64% at Y1 to 80% by Y5, though there was still considerable attrition from, and switching between Ac Research, Fp Research, and G/Np Research during the first five years after graduation (Fig. 5). Taken together, these comparisons show that alumni who pursue Teaching and AMO careers immediately after PhD completion are more likely to persist in the identical career at Y10 than alumni who enter research career areas.

**Table 10.** Percent of alumni at 1, 3, or 5-years post-PhD whose career area is identical to their career area at Y10.

|  | Years post-PhD |  |  | Career area at Y10 |
| --- | --- | --- | --- | --- |
|  | One | Three | Five |  |
| **Percent of alumni working in an identical career area as ten years post-PhD** | 42% | 54% | 69% | *All Career Areas* |
|  | 64% | 71% | 80% | *Some Type of Research* |
|  | 41% | 52% | 67% | Ac Research |
|  | 58% | 54% | 63% | Fp Research |
|  | 12% | 20% | 38% | G/Np Research |
|  | 80% | 83% | 89% | Teaching |
|  | 98% | 97% | 94% | AMO |

Alumni who graduated 1/1/1997 to 12/31/2011 were included in this analysis (n=654; Cohort E). The first row shows the percentage of alumni whose career area at Y10 matched their career area at an earlier timepoint across all career areas, regardless of the specific career area of employment. The second row shows the percentage of alumni with Some Type of Research career at Y10 as well as the earlier timepoint, but not necessarily the same Research career area. The raw number of alumni underlying each percentage calculation is shown in S3 Table.

### Evolution of early career relative to career goal and first non-training position

For the subset of alumni for whom we had an exit survey and a Y10 career outcome (Cohort F: 271 alumni who graduated between 2005-2011), we analyzed whether their career goal at defense matched their Y10 career area. Except for AMO careers, the distribution of career outcomes at Y10 appeared comparable to the distribution of career goals at defense (Fig. 6). However, across all career areas, only 51% of students’ career goals at defense matched their career outcome at Y10 (Table 11). Alignment of career goal at defense with career area at Y10 was highest for those with AMO or Ac Research career goals (80% and 52%, respectively) and lowest for those interested in G/Np Research careers (13%). At Y10, more than half the 27 individuals who were Und/s at defense were in some kind of Research career (Table 11).

**Fig 6.**
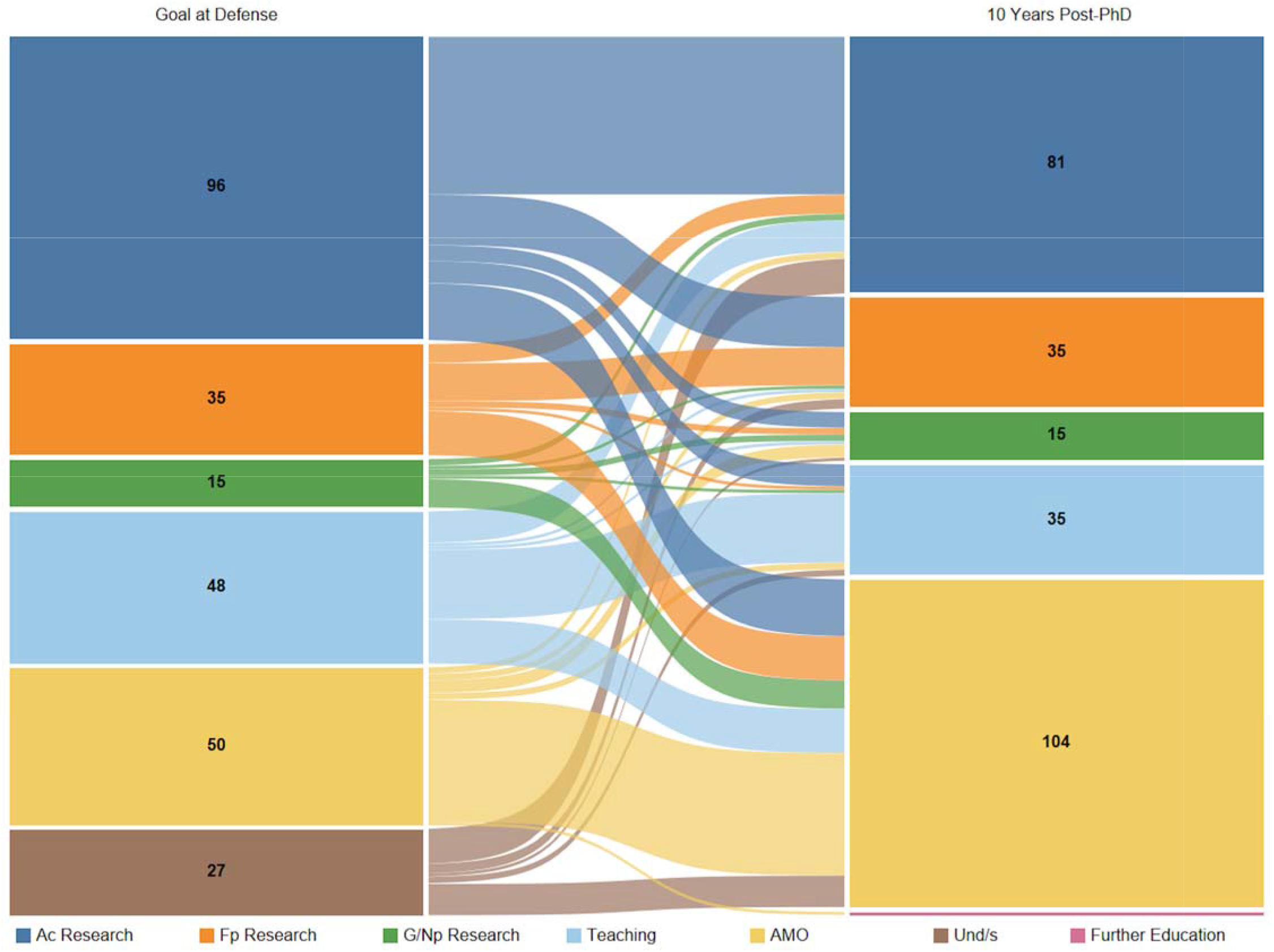
Comparison of career goal at defense to career area at 10 years after graduation. Sankey diagram showing career goals of biomedical sciences PhD students at defense (left nodes) compared to career area at 10 years after graduation (right nodes). The color of the links corresponds to the students’ career goal at defense and the thickness of each link is proportional to the number of alumni represented in the link. Students who graduated between 1/1/2005 and 12/31/2011 and completed the exit survey were included in this analysis (n=271, cohort F).

**Table 11.**
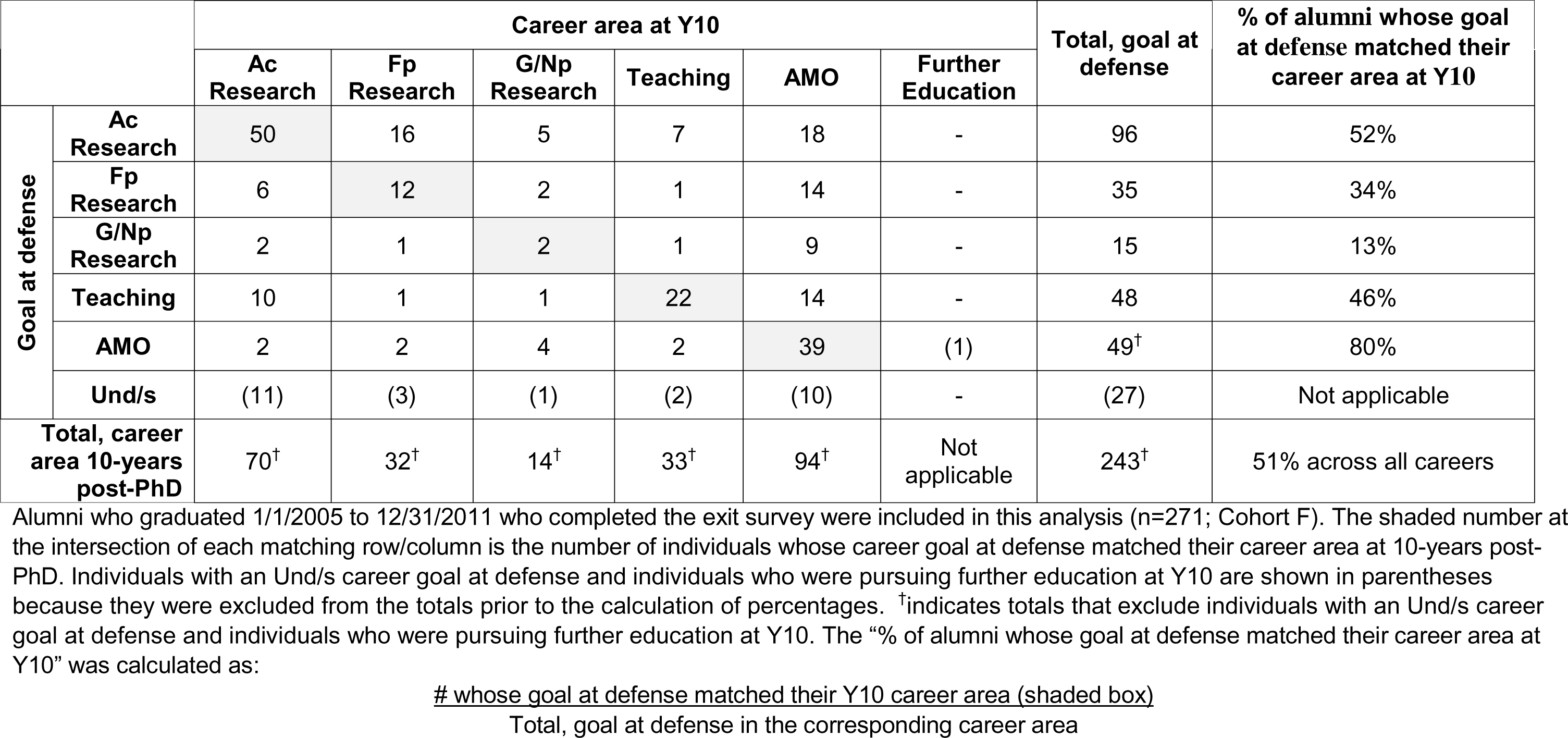
Goal at defense compared to career area at Y10.

|  |  | Career area at Y10 |  |  |  |  |  | Total, goal at defense | % of alumni whose goal at defense matched their career area at Y10 |
| --- | --- | --- | --- | --- | --- | --- | --- | --- | --- |
|  |  | Ac Research | Fp Research | G/Np Research | Teaching | AMO | Further Education |  |  |
| Goal at defense | Ac Research | 50 | 16 | 5 | 7 | 18 | - | 96 | 52% |
|  | Fp Research | 6 | 12 | 2 | 1 | 14 | - | 35 | 34% |
|  | G/Np Research | 2 | 1 | 2 | 1 | 9 | - | 15 | 13% |
|  | Teaching | 10 | 1 | 1 | 22 | 14 | - | 48 | 46% |
|  | AMO | 2 | 2 | 4 | 2 | 39 | (1) | 49<sup>†</sup> | 80% |
|  | Und/s | (11) | (3) | (1) | (2) | (10) | - | (27) | Not applicable |
| Total, career area 10-years post-PhD |  | 70<sup>†</sup> | 32<sup>†</sup> | 14<sup>†</sup> | 33<sup>†</sup> | 94<sup>†</sup> | Not applicable | 243<sup>†</sup> | 51% across all careers |

Alumni who graduated 1/1/2005 to 12/31/2011 who completed the exit survey were included in this analysis (n=271; Cohort F). The shaded number at the intersection of each matching row/column is the number of individuals whose career goal at defense matched their career area at 10-years post-PhD. Individuals with an Und/s career goal at defense and individuals who were pursuing further education at Y10 are shown in parentheses because they were excluded from the totals prior to the calculation of percentages. <sup>†</sup>indicates totals that exclude individuals with an Und/s career goal at defense and individuals who were pursuing further education at Y10. The “% of alumni whose goal at defense matched their career area at Y10” was calculated as:

We also examined the match between the career area of alumni first non-training position and their Y10 job (Fig. 7 and Table 12). There were 669 alumni who graduated from 1997-2011 for whom we had both their first non-training position milestone and their career outcome at the Y10 timepoint. Seven of these alumni were in Further Education at Y10 so they were excluded from the analysis.

**Fig 7.**
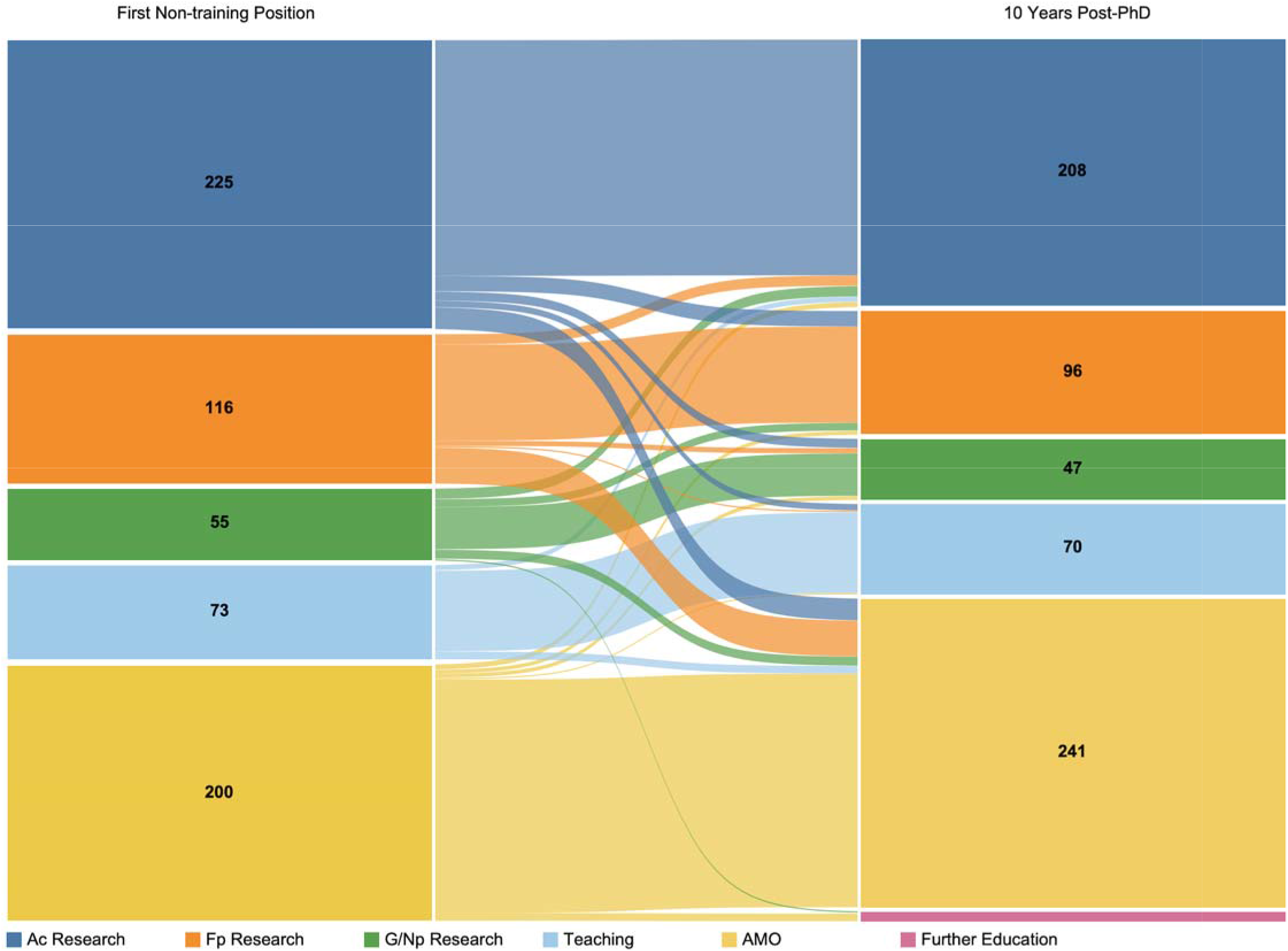
Comparison of first non-training position to career area at 10 years after graduation. Sankey diagram showing the first non-training position of biomedical sciences PhD alumni (left nodes) compared to career area at 10 years after graduation (right nodes). The color of the links corresponds to the career area of the first non-training position and the thickness of each link is proportional to the number of alumni represented in the link. Students who graduated between 1/1/1997 and 12/31/2011 were included in this analysis (n=669, cohort G).

**Table 12:** First non-training position compared to career area at Y10.

|  |  | Career area at Y10 |  |  |  |  |  | Total, 1<sup>st</sup> non-training position | % of alumni with matching career area in the 1<sup>st</sup> non-training position and Y10 |
| --- | --- | --- | --- | --- | --- | --- | --- | --- | --- |
|  |  | Ac Research | Fp Research | G/Np Research | Teaching | AMO | Further Education |  |  |
| First non-training position | Ac Research | 184 | 12 | 7 | 5 | 17 | - | 225 | 82% |
|  | Fp Research | 8 | 75 | 4 | 1 | 28 | - | 116 | 65% |
|  | G/Np Research | 8 | 6 | 33 |  | 7 | (1) | 54<sup>†</sup> | 61% |
|  | Teaching | 4 |  |  | 63 | 6 | - | 73 | 86% |
|  | AMO | 4 | 3 | 3 | 1 | 183 | (6) | 194<sup>†</sup> | 94% |
| Total, career area at Y10 |  | 208<sup>†</sup> | 96<sup>†</sup> | 47<sup>†</sup> | 70<sup>†</sup> | 241<sup>†</sup> | (7) | 662<sup>†</sup> | 81% across all career areas |

Alumni who graduated 1/1/1997 to 12/31/2011 for whom we had a known first non-training position and Y10 career were included in this analysis (n=669; Cohort G). The shaded number at the intersection of each matching row/column is the number of alumni with matching career areas in the first non-training position and at Y10. Individuals who were pursuing further education or training at Y10 are shown in parentheses because they were excluded from the totals prior to the calculation of percentages. <sup>†</sup>indicates totals that excluded individuals pursuing further education or training at Y10. The “% of alumni with matching career area in the 1<sup>st</sup> non-training position and Y10” was calculated as:

In the remaining cohort, the relative distribution of first non-training positions was very similar to the distribution of alumni employment at Y10 (Fig. 7). There was a 1-3% loss of alumni employed in Ac Research, Fp Research, G/Np Research, and Teaching career areas between the first non-training position and Y10, which led to a net growth of alumni employed in AMO careers at Y10. Notably, once alumni acquired their first non-training position, persistence in the same career area at Y10 was 81%, and persistence was particularly high for alumni whose first non-training position was an AMO career or a career in Ac Research or Teaching (Table 12). Alumni who moved out of a first non-training position in Ac Research primarily moved to either Fp Research or AMO jobs at Y10, and those who moved out of a first non-training position in Fp Research mainly moved to AMO jobs. Those who moved out of G/Np Research to another career type by Y10 diverged evenly to Ac Research, Fp Research, or AMO careers; none changed to Teaching. Persistence in the same career area at Y10 was highest (94%) among those whose first non-training position was an AMO job, and the few who moved out of AMO primarily moved into some type of Research job. Teaching was also highly stable, and those who moved to another career type by Y10 moved into either Ac Research or AMO careers.

## Discussion

Here, we present the results of a longitudinal study that links the career goals of biomedical PhD students at matriculation and defense to their career outcomes at key career milestones and timepoints up to ten years after completing PhD training. Although our study was restricted to 1,452 alumni from a single institution, the demographics of our student population are similar to that of other institutions ^12, 13^ and our students’ time to degree and total publication number are similar to those at other BEST institutions ^14^, suggesting that our findings may be generalizable. Several groups have reported on either PhD student career goals ^12, 15–19^ or career trajectories after PhD completion ^13, 20–22^, but to our knowledge, this is the first study to examine the two in combination. This type of analysis is rare because it requires longitudinal data collection for an extended period as well as an in-depth understanding of science-related careers to curate alumni career outcomes accurately. However, we hope that other institutions will be inspired to track and report similar career goal and milestone data, thus enabling cross-institutional analyses to detect trends in the early careers of biomedical research scientists.

Our career outcomes taxonomy (S1 Appendix) expands on previous classification systems ^9, 10^ with the addition of flags to document faculty positions as well as the first position after PhD completion and the first non-training position of PhD graduates. The use of these novel flags enabled us to report on career milestones and better understand the early career trajectories of our alumni. Major findings include that nearly equal numbers of alumni were in Ac Research and AMO careers at Y10, and the best indicator of alumni career area 10 years after graduation with a PhD was their first non-training position, whether that occurred immediately after PhD completion or after a period of postdoctoral training.

### Changing career goals

More than half the PhD students in our study population changed their primary long-term career goal between matriculation and defense. Others have also observed that a large fraction of graduate students change career goals during graduate school ^12, 15–17, 19, 21, 23, 24^, indicating that this is common behavior rather than the exception to the rule.

As reported by Wood et. al. (2020) ^17^, we found that students who were interested in research-intensive faculty careers at defense were a mix of those who were interested in PI careers since matriculation and those who became interested in being a PI during graduate training. Nonetheless, in contrast to the Wood study, we observed a 19% net decrease in the number of students interested in research-intensive faculty careers during PhD training, a trend that is consistent with most other studies ^12, 15, 16, 19, 21, 23^. The reasons cited by our students for changing career goals during graduate school fit well within the framework of Social Cognitive Career Theory (SCCT) ^25^. SCCT posits that an individual’s career interests are formed through a dynamic interplay between self-efficacy in a particular domain and outcomes expectations about careers in that area. In our analysis, the internal factors cited by students as reasons for changing career goals contribute to formation of self-efficacy, and the external factors cited by students as reasons for changing career goals align with their outcomes expectations about attaining a career as a PI. Several studies designed to examine career interest formation through the lens of SCCT have also shown that social identity influences career interests of biomedical scientists ^15, 19, 23, 26, 27^, though we did not attempt to analyze the role of social identity in our population, as it is beyond the scope of this study.

Changes in career interest also occur during postdoctoral training, as we found that only 51% of our alumni had a first non-training position in a career area that matched their career goal at defense. We did not survey our alumni directly to ask why they changed directions during postdoctoral training, but a recent survey study by Lambert and colleagues (2020) ^26^ suggested that success in research during postdoctoral training played an important role in shaping a postdoc’s self-efficacy and intent to pursue a research-intensive faculty career. This large survey of postdocs from over 80 institutions also found that social identity, personal values, and perceptions about faculty careers were significant factors in determining postdocs’ career interests.

The fluidity of PhD student career interests during graduate school, and the subsequent shift in career trajectory that occurs yet again during postdoctoral training, points to the critical need for research mentors and training programs to provide both student and postdoc trainees with career mentoring and encourage experiential learning about research- and research-related careers, including that of academic scientists. Historically, many institutions and faculty did not promote career development activities for trainees because of perceived conflict of effort with the NIH funding sources from which many biomedical trainee stipends are paid ^28^. However, shortly after the NIH created the NIH Director’s Broadening Experiences in Scientific Training (BEST) program, which funded 17 institutions to test and evaluate the impact of providing grad students and postdocs with experiential training for a range of research- and research-related careers, the NIH clarified that it is acceptable for PhD students and postdocs supported by NIH grants to participate in career development activities ^29^.

Other faculty were historically reluctant to encourage trainees to participate in career development activities because they believed that spending time on career development activities would diminish trainee productivity and increase time in training. These concerns were debunked by a robust cross-institutional analysis from ten NIH BEST institutions, who pooled their data on student participation in career development activities to demonstrate conclusively that engaging in such activities did not extend time-to-degree in graduate school or decrease the number of first-author or total publications of graduate students ^14^. Furthermore, the NIH BEST experiment demonstrated that there are a variety of ways for PhD students and postdocs to engage in meaningful experiential career learning opportunities in non-academic laboratory work environments, many of which do not require extended time away from the laboratory ^30–36^.

### Most biomedical PhD alumni did a postdoc, regardless of career goal

Our data show that between 1997-2021, the majority of our biomedical sciences PhD graduates entered a postdoctoral position, primarily in Ac Research. During postdoctoral training, biomedical scientists build out their CV, expand their scientific expertise, and develop autonomy that is valuable preparation for research-intensive careers, and especially for careers as PIs who oversee an independent research program ^37, 38^. Postdoctoral training is also considered an asset for many teaching-intensive colleges that expect faculty to engage undergraduates in laboratory research.

Given the benefits of postdoctoral training, it would be easy to assume that alumni who do postdocs intend to pursue a research- or teaching-intensive faculty position that involves significant research. Indeed, nearly 90% of our students with a Research career goal at defense, and nearly 75% with a Teaching career goal, pursued postdoctoral training. Furthermore, alumni with a Research career goal who did a postdoc were more likely to enter a Research career for their first non-training position than those who did not do a postdoc (84% compared to 52%).

However, the 76% of our alumni who did a postdoc far exceeded the 31% who had an Ac Research career goal at defense, a finding consistent with previous studies and reports ^37, 39^. This imbalance was partly because 53% of individuals with an AMO career goal and 74% of those who were Und/s at defense chose to do postdocs. Our alumni with an AMO career goal were more likely to attain a first non-training position quickly compared to alumni with an Ac Research career goal, but economic analyses have shown that time spent in postdoctoral training has negative financial ramifications, especially for individuals who do not pursue Ac Research careers ^20, 38, 40, 41^. Although postdoctoral training may be beneficial experience for some AMO positions (e.g., editorial careers at scientific journals), it is not a requirement for many AMO jobs. We did observe that alumni in our study who were in AMO or Teaching careers at Y10 spent less time in postdoctoral training than individuals who pursued Research careers, but the opportunity cost of doing a postdoc on lifetime earnings should prompt institutions to help PhD students who do not intend to pursue Research careers to move quickly into their first non-training position, skipping the postdoctoral training period if it isn’t necessary or of benefit for their intended career.

### The pursuit of postdoctoral training declined over time

We observed that the percentage of alumni who pursued postdoctoral training has declined steadily over time, dropping to below 60% for the first time in 2019. Our institutional data mirrors national trends in postdoctoral employment that have been the focus of several recent articles ^42–44^. Concern about declining numbers of postdocs is the basis for the creation of a new Advisory Committee to the NIH Director that is charged with providing recommendations to address the causes and consequences of declining numbers of postdocs ^11^.

Most of our PhD students who did not do a postdoc immediately after PhD completion entered the career area that matched their goal at defense, suggesting that postdoctoral training was not necessary for them to attain their preferred career. The recent trend toward fewer of our PhD alumni pursuing postdoctoral training may be an encouraging sign that PhD students are defining their career goals earlier in their scientific training rather than deferring career decision-making and pursuing postdoctoral training by default ^37^. If so, interventions like the NIH BEST program, which were designed to broaden PhD and postdoc training and perspectives to better align with the realities of the biomedical research workforce, appear to be working. The resulting decline in postdocs is an understandable cause for concern among academic institutions that have relied upon postdoctoral scientists and early career researchers to make biomedical research discoveries. However, as noted in a recent editorial by Dzirasa (2023) ^45^, the solution to this problem requires robust policy responses and economic investment aimed at making academic research careers more sustainable and attractive.

### Evolution of early career pathways

Following individual careers longitudinally for the first ten years post-PhD revealed that our alumni gradually moved out of Ac Research and into other career areas. This shift over time is consistent with data from a study of Wayne State University STEM alumni ^13^. An important distinction is that we followed career outcomes for the same individuals over time, whereas the Mathur study reported career outcomes for three different groups of alumni, at three distinct timepoints. Following the same individuals over time allowed us to determine that the shift of our alumni out of the Ac Research career area in the first ten years after graduation was largely due to alumni finishing postdoctoral training. Within 10 years of earning their doctoral degree, our alumni were employed in Ac Research or Teaching and held tenure-track faculty jobs at rates comparable to biomedical PhDs nationally ^4^.

We were particularly interested in comparing patterns of alumni employment ten years after graduation to earlier timepoints and milestones to determine the point at which alumni become relatively set in their career path. Not surprisingly, only 51% of alumni were employed in a career area at Y10 that matched their career goal at defense. While an increasing percentage of alumni at Y1, Y3, and Y5 were employed in the same career area as they were employed at Y10, the continued movement of alumni out of postdoc positions meant that none of these earlier timepoints were particularly good indicators of the career area of employment of alumni at Y10. It is important to note that except for the transition out of postdoctoral training, our analyses focused on the movement of alumni between different career areas, not between jobs per se. For example, an alumnus who changed from being a medical writer at Y3 to being a medical science liaison at Y5 would be represented in the AMO career area at both timepoints. Thus, our visualizations do not depict the number of job changes alumni had in the first ten years after graduation.

Interestingly, the career area that alumni entered for their first non-training position was identical to their career area at Y10 for 81% of alumni, suggesting that this milestone is a good indicator of the early career pathway of alumni. This observation has several implications for policymakers, trainees, and research training programs. Currently, the NIH requires institutions with T32 training grants to track the career outcomes and publications of PhD alumni for 15 years after degree conferral. The amount of data collection and reporting that is required for T32 training grant submissions is time-consuming and represents a substantial administrative burden. Our findings suggest that alumni could be tracked to their first non-training position, whenever that occurs, as a reasonable proxy for the broad career area in which they will likely be employed in the future. Training programs could continue to collect the publication record of alumni who pursue Research careers for the purpose of understanding the impact of the T32 training program on scientific research advances, but the administrative burden of tracking alumni careers beyond their first non-training position could be eliminated. Many institutions will undoubtedly continue tracking alumni for other purposes with a frequency that makes sense for their internal needs.

For trainees, it is interesting to note that the career area in which one works during their first job outside of training – whether it occurs after graduate school or after a postdoc – is likely to be the career area in which one will work at Y10; this is especially true if one’s first step out of training is an AMO or Teaching career. While there is a decreasing trend toward pursuing postdoctoral training, PhD students who are interested in a Research career should strongly consider postdoctoral training, as we observed that alumni whose primary long-term career goal at defense was a Research career, but who did not do a postdoc, were less likely to attain a Research career for their first non-training position. Conversely, trainees who have ruled out careers in Research should be actively supported by their institutions and training programs to skip postdoctoral training and pursue other career options that utilize their transferable skills, biomedical sciences expertise, and research training in ways that better align with their interests. Doctoral training provides scientists with many transferable skills that are valued by PhD alumni and employers in a range of professions ^46, 47^, and PhD scientists contribute to the biomedically trained workforce in myriad productive ways, both in research and research-related careers. Training programs should aim to help connect trainees to these careers quickly to maximize their satisfaction and contributions to the scientific enterprise and keep graduate school an attractive path for high potential individuals.

## Materials and Methods

### Study population

The population examined in this paper included graduates of Vanderbilt University’s biomedical sciences PhD programs who (a) were admitted in 1992 or after through one of two umbrella PhD admissions programs, the Interdisciplinary Graduate Program (IGP) or the Quantitative and Chemical Biology Program (QCB), or (b) were directly admitted or transferred to a program affiliated with IGP/QCB programs and (c) graduated with a PhD between 1997 and August 31, 2021. Data from six deceased individuals was excluded from all datasets examined. Students who were enrolled in Vanderbilt’s dual MD/PhD program were not part of the study population as their career outcomes 10 years after earning a PhD can be very different from those of the population reported in this paper. Sixty-nine alumni were excluded from all analyses because they did not have an exit survey, a first known position after PhD, and a known position for at least one of the timepoints examined in this analysis (1, 3, 5, or 10 years after PhD).

Of the 1,452 PhD alumni who graduated during this time period, 99% earned a PhD from one of the following current or legacy biomedical PhD programs: Biochemistry, Biological Sciences, Cancer Biology, Cell Biology, Cell and Developmental Biology, Chemical and Physical Biology, Human Genetics, Microbe-Host Interactions, Microbiology and Immunology, Molecular Biology, Molecular Pathology and Immunology, Molecular Physiology and Biophysics, Neuroscience, or Pharmacology; the remaining 1% of students earned a PhD in Biomedical Engineering, Biomedical Informatics, Chemical Engineering, Chemistry, Interdisciplinary, Physics, or Psychology.. The *n* for analyses varied based on the timepoint or milestone restrictions for the analysis and the maximum cohort size for any figure was n=1,413. Table 1 shows the inclusion/exclusion criteria for each figure. All cohorts analyzed had the following demographic breakdown: 52-60% female, 8-15% underrepresented in the biomedical sciences, and 14-19% foreign nationals with a temporary US visa, and no demographic group was over- or underrepresented relative to our population of biomedical sciences PhD students at Vanderbilt.

### Exit survey

Starting on January 1, 2007, all biomedical PhD students at Vanderbilt University completed a voluntary online exit survey (IRB #171216) within 2 to 3 weeks after their dissertation defense. The exit survey was implemented to collect student feedback on their PhD training, career goals, and immediate employment plans. A total of 944 students completed the exit survey, including twenty alumni who defended in 2005 and 2006 and served as a focus group to help validate the survey. The overall survey response rate from 1/1/2007 to 8/9/2021 was 90%, calculated as the [(number of students who completed the survey) ÷ (# of students invited to complete the survey)].

In the exit survey, students were asked the following questions about their career goals at matriculation and defense: (1) “Which best describes your primary long-term career goal when you STARTED graduate school?” and (2) “Which best describes your primary long-term career goal NOW, as you finish graduate school?”. Answer options for both questions were: Faculty position in academia (primarily research), Faculty position in academia (primarily teaching), Science Outreach/K-12 Education, Research position in industry, Research position in government/nonprofit lab, Other (please specify); “I didn’t know what I wanted to do after graduate school” was also an answer option for the question about career goals at matriculation. In 2009, the following answer options were added: Science or Medical Writing/Editing/Publishing, Patent Law, Science Policy, and Undecided. The exit survey also contains an optional, free-text question to gather information about why students changed their career goals during graduate school: “If your career goals changed since you started your PhD program, what factor has most affected these goals?” While asking students to report their career goal from several years earlier may have the potential for recall bias, our results on changing career goals during PhD training are consistent with previous studies that examined student career goals at the specific timepoints in question ^16, 21^ as well as studies that asked students to recall their career goals from earlier timepoints^12, 15, 23^.

### Coding of free-text responses

We analyzed free-text responses about reasons for changing career goals from two groups: those whose career goals between matriculation and defense changed (1) *away from* or (2) *toward* becoming a research-intensive faculty member in academia. Out of 944 exit survey respondents, there were 171 individuals in the first category, 153 of whom provided comments. There were 100 individuals in the second category, 68 of whom provided reasons for switching career goals.

The coding process began with a member of the research team using inductive thematic analysis to identify semantic themes in the comments of both groups ^48^. In a first reading of comments, two main types of themes were identified that we called either: (1) external factors, relating to the respondents’ perceptions about the job of being a PI or working in academia and (2) internal factors, relating to the respondents’ perceptions about themselves or fit with a career option, or their personal experience in graduate school. In a second reading of comments, eight sub-themes (four external and four internal) were noted for individuals whose career goals changed away from being a research-intensive faculty member in academia, and four sub-themes (all internal) were identified for individuals whose career goals changed toward being a research-intensive faculty member in academia. To be defined as a sub-theme, a sentiment’s prevalence across the data set was considered, and a sentiment had to be expressed by at least ten people within either group of survey respondents. Themes and sub-themes were verified by a second member of the analysis team, then both team members independently coded each comment. A single comment could be assigned multiple codes if the survey respondent mentioned more than one theme or sub-theme in their survey response. The two coders resolved discrepancies in coding by clarifying sub-theme definitions and consensus building until there was complete agreement in coding. Nonspecific responses for both groups were coded, “Not descriptive enough to categorize.”

### Career outcomes

The career outcomes of biomedical PhD alumni were collected using Google searches of publicly available information, including PubMed, ResearchGate, ORCID, LinkedIn, and institutional websites. Alumni who could not be found by a single member of our team were included in follow-up searches by two additional team members. Specific data captured for each position held by an alumnus included job title, employer, location of employment, and inferred start date. Position start dates allowed us to determine the length of time alumni remained in each position by calculating the time from the start date of the first position to the start date of the second position. We were able to find career outcomes for 95% of our alumni who graduated between January 1, 1997, and August 31, 2021. Employment information was stored in a database we developed and optimized for tracking employment outcomes over time.

### Relational database and data processing

A PostgreSQL relational database was created to store static academic records and dynamic outcomes data about our graduates. The database is deployed onto a secure institutional server, provides multi-user access, and maintains project data integrity. To generate reports and visualize the data, we used commercially available software tools, Aqua Data Studio and Tableau; database revision history and process documentation are stored in a wiki, Atlassian.

### Career categorization

To categorize alumni careers, we used a three-tiered taxonomy developed in-house in 2012 to classify career outcomes based on sector of employment, type of career, and job function (S1 Appendix). Our strategy to classify each position by sector and career activity was based on the categorization of STEM PhD careers in the NSF Survey of Doctoral Recipients. Our initial list of job functions was developed organically by grouping similar careers together, then subsequently modified to achieve greater consistency with the classifications used in cross-institutional surveys by the BEST consortium ^49^. Between 2012-2015, we refined our categorization system to include an additional taxonomy for classifying faculty jobs, capturing secondary jobs (e.g., adjunct professorships or starting a side-business; only primary positions were included in this study), flagging career milestones of the first position after PhD completion and the first non-training position, assigning Carnegie IDs to classify employment in US institutions of higher education, and flagging positions that are science-related, entrepreneurial, or executive-level

The classification system we developed is similar to the three-tiered Unified Career Outcomes Taxonomy ^9^ and further optimized by Stayart et. al. (2020) ^10^, with differences described in S1 Appendix. At least two experienced career outcomes professionals independently categorized every job, and every position that was not identically classified was reviewed to reach a consensus category.

For the analyses in this paper, we also used our taxonomy to categorize student career goals at the time of matriculation and defense. Respondents who answered, “Other (please specify)” were asked to describe their career goals in free text, and each of these responses was coded to the best of our ability. Some “Other” responses were very specific and easy to code in all three tiers of our taxonomy, e.g., “lab manager in academia” was coded as Career Sector: “Academia”, Career Type: “Primarily Research”, Job Function: “Research Staff or Technical Director”. In contrast, some “Other” responses were nonspecific and could not be coded in all three tiers of the taxonomy. In these cases, we coded the available tier(s) and coded the remaining tier(s) as, “Unspecified.” For example, an “Other” response described as, “Industry” was coded as Career Sector: “For-profit”, Career Type: “Unspecified” and Job Function “Unspecified”. Free-text responses that indicated that the student was unsure (e.g., “I don’t know”) or considering multiple options (e.g., "science writing or clinical research”) were coded as, “Undecided.”

### Timepoints and milestones

This study examined alumni career outcomes at four timepoints after graduation, and at two key milestones that are important for understanding the career trajectories of biomedical scientists: the first position after completion of the PhD and the first non-training position. The career outcome at 1, 3, 5 or 10 years after graduation was defined as the job held by an alumnus on the exact date 1, 3, 5, or 10 years after the date of degree conferral recorded by the Vanderbilt Graduate School; Vanderbilt University has three standard graduation dates annually, in May, August, and December. The two milestones were manually designated (or “flagged”) by career outcomes coders. The first non-training position was defined as the first job that is not postdoctoral training, a degree-granting program, or some other term-limited training position (e.g., medical residency or fellowship.)

The first position after completion of the PhD was defined as the first job or postdoctoral position that is *not* in the same lab in which the alumnus completed their PhD research. An exception to this rule was made if a postdoctoral position in the dissertation lab lasted longer than 6 months. We developed this procedure because about 30% of our alumni did a short postdoc of 6 months or less in their dissertation lab prior to taking a job or full-term postdoc in another lab. Alumni reported in the exit survey that they did these short postdocs for a variety of reasons, often to time a move to coincide with the start date of their next position or to coincide with the needs of their family or partner. Although short postdocs in a PhD lab were not flagged as the first position after PhD completion unless they were longer than 6 months, the duration of these short postdocs was included in the calculation of time to first non-training position and duration of postdoctoral training.

### Statistics

Logistic regression was used to examine the significance of trends in alumni entering postdoctoral training as their first position after PhD completion. For regression analysis, graduation year was treated as the independent variable, and choice to enter postdoctoral training as a binary (yes/no) dependent variable. Regression analysis and visualization, likelihood ratio testing, and calculation of odds ratios were conducted in R v. 4.2.2 ^50^ using lmtest ^51^ and ggplot2 ^52^ packages. Percent decrease in probability was calculated as the difference in probabilities of entering a postdoctoral position at the initial full-year timepoint (1997) and end full-year timepoint (2020), all divided by the probability at the initial timepoint.

For alumni who entered at least one postdoctoral training position, the length of time between a graduate’s doctoral thesis defense and entry into their first non-training position was calculated. These calculated time-to-event values were used to generate cumulative incidence curves grouped by individuals’ career goals at defense, which represent an individual’s probability of entering a first non-training position at any point in time following doctoral thesis defense. Cumulative incidence curves were generated using 1 - survival probability determined by the Kaplan-Meier survival calculation method ^53^. This method is robust in that it allows for measurements from individuals who have not entered a first non-training position by the end of the observation period ("censored" individuals) to contribute to calculations for the probability of entering a first non-training position. Observations used in Kaplan-Meier calculations were coded into two separate groups: 1) non-censored (observations from alumni who entered their first non-training position), and 2) censored (observations from alumni who defended their doctoral thesis but had not entered their first non-training position by 5/1/2020, the cut-off date for recorded observations). For non-censored observations, time-to-event was measured as the difference between doctoral thesis defense and entry into first non-training position. For censored observations, time-to-event was measured as the difference between doctoral thesis defense and the cut-off date for recorded observations. One individual pursued a first non-training position prior to defending their thesis, resulting in a negative time in training calculation; this individual was not included in the analysis. Kaplan-Meier calculations make use of both censored and non-censored observations, providing probability inputs for the final cumulative incidence curve. Probability calculations and visualizations for cumulative incidence curves were conducted using GraphPad Prism v. 9.5.1 for Windows (GraphPad Software, San Diego, California USA).

For alumni with known career outcomes 10 years after graduation, Kruskal-Wallis with Dunn’s multiple comparisons testing was conducted on data reflecting graduates’ total time spent in active postdoctoral training. Statistical significance was determined using an alpha threshold of 0.05. Both analyses were conducted using GraphPad Prism.

### Data visualization

To visualize how career goals and career outcomes evolved over time, the Sector and Career Type tiers for research-focused careers were grouped as either Academic Research (Ac Research), For-profit Research (Fp Research), or Government or Nonprofit Research (G/Np Research). Non-research careers in all sectors were grouped together and presented as either Teaching careers or Administrative, Managerial, or Operational (AMO). Examples of job titles held by individuals in these other career areas are shown in S2 Table. Students with Undecided or Unspecified career goals at defense were grouped together for visualization as “Und/s.”

### Not employed, Unknown, and Not Science-Related

Some alumni are hard to locate, for a variety of reasons. In instances where one team member could not locate an alumnus, two other people attempted a search and verified that the individual could not be found. During the time period covered in this study (1997-2021), ten alumni were not employed, and 88 alumni had unknown positions at one or more of the defined milestones or timepoints of interest (first position after PhD, first non-training position, or 1, 3, 5, or 10 years after graduation with a PhD). These individuals were included in the figures in which they had known positions and excluded from figures and data analyses in which they had a missing milestone or timepoint. Details of the inclusion/exclusion criteria for cohorts represented in each figure and table are described in Table 1. The 26 alumni (1.7% of our population) who held one or more jobs that were flagged as “not science-related” were included in the visualizations.

## Author Contributions

Abigail M. Brown - Conceptualization, Data curation, Investigation, Methodology, Project administration, Resources, Validation, Writing – original draft

Lindsay C. Meyers - Data curation, Formal Analysis, Investigation, Methodology, Project administration, Resources, Software, Validation, Visualization, Writing – review & editing

Janani Varadarajan - Data curation, Formal Analysis, Investigation, Methodology, Project administration, Resources, Software, Validation, Visualization, Writing – review & editing

Nicholas J. Ward – Formal Analysis, Software, Validation, Visualization, Writing – review & editing Jean-Philippe Cartailler – Resources, Software

Roger G. Chalkley – Funding Acquisition, Writing – review & editing

Kathleen L. Gould – Funding Acquisition, Supervision, Writing – review & editing

Kimberly A. Petrie - Conceptualization, Data curation, Formal Analysis, Funding Acquisition, Investigation, Methodology, Project administration, Resources, Supervision, Validation, Visualization, Writing – original draft

## Data Availability Statement

The data that support the findings of this study are openly available in OSF at https://osf.io/f28vb/.

## Acknowledgements

We would like to thank the Vanderbilt Biostatistics Department for providing guidance on time-to-event and trend analyses for alumni who entered postdoctoral training. We are grateful to former interns Rachel Howell, Nick Catalan, Chiedza Chauruka, and Rian Djita for their assistance in collecting career outcomes for our alumni. The Sankey diagrams were made using the Multi-Level Sankey template created by Ken Flerlage with credits to Jeff Shaffer and Olivier Catherin for original development (www.moxyanalytics.com). Finally, we are grateful to Madhvi Venkatesh for her valuable insights and thoughtful suggestions before submission.

## Abbreviations

Ac Research: Academic Research;
AMO: Administrative, Managerial, or Operational;
BEST: Broadening Experiences in Scientific Training;
CI: Confidence Interval;
Fp Research: For-profit Research;
G/Np Research: Government or Nonprofit Research;
IGP: Interdisciplinary Graduate Program;
NIH: National Institutes of Health;
NRSA: National Research Service Awards;
NSF: National Science Foundation;
ORCID: Open Researcher and Contributor Identifier;
PI: Principal Investigator;
QCB: Quantitative and Chemical Biology Program;
SCCT: Social Cognitive Career Theory;
STEM: Science Technology Engineering and Mathematics;
Und/s: Undecided or Unspecified;
Y1: one year after graduation with a PhD;
Y3: three years after graduation with a PhD;
Y5: five years after graduation with a PhD;
Y10: ten years after graduation with a PhD

## References

1. *National Research Council. Personnel Needs and Training for Biomedical and Behavioral Research:* 1983 Report. 1983:232. https://nap.nationalacademies.org/catalog/19456/personnel-needs-and-training-for-biomedical-and-behavioral-research-the

2. Garrison HH, Gerbi SA. Education and employment patterns of U.S. Ph.D.’s in the biomedical sciences. FASEB J. Feb 1998;12(2):139–48. doi:10.1096/fasebj.12.2.139

3. National Center for Science and Engineering Statistics (NCSES). Survey of Doctorate Recipients, 2021. NSF 23-319. National Science Foundation. https://ncses.nsf.gov/pubs/nsf2319

4. Biomedical Workforce Working Group Report of the Advisory Committee to the Director. National Institutes of Health. Accessed February 26, 2023, https://acd.od.nih.gov/documents/reports/Biomedical_research_wgreport.pdf

5. McDowell GS, Gunsalus KT, MacKellar DC, et al. Shaping the Future of Research: a perspective from junior scientists. F1000Res. 2014;3:291. doi:10.12688/f1000research.5878.2

6. Alberts B, Kirschner MW, Tilghman S, Varmus H. Opinion: Addressing systemic problems in the biomedical research enterprise. Proc Natl Acad Sci U S A. Feb 17 2015;112(7):1912–3. doi:10.1073/pnas.1500969112

7. Hitchcock P, Mathur A, Bennett J, et al. The future of graduate and postdoctoral training in the biosciences. Elife. Oct 19 2017;6doi:10.7554/eLife.32715

8. Cheng SD. Where Are All the Scientists? Resources for Studying the Long-Term Careers of STEM PhDs. White paper for National Bureau of Labor Statistics. 2020. https://www.nber.org/programs-projects/projects-and-centers/value-medical-research/value-medical-research

9. Mathur A, Brandt P, Chalkley R, et al. Evolution of a Functional Taxonomy of Career Pathways for Biomedical Trainees. J Clin Transl Sci. Apr 2018;2(2):63–65. doi:10.1017/cts.2018.22

10. Stayart CA, Brandt PD, Brown AM, et al. Applying inter-rater reliability to improve consistency in classifying PhD career outcomes. F1000Res. 2020;9:8. doi:10.12688/f1000research.21046.2

11. National Institutes of Health. ACD Working Group on Re-envisioning NIH-Supported Postdoctoral Training. Accessed February 26, 2023, https://acd.od.nih.gov/working-groups/postdocs.html

12. Fuhrmann CN, Halme DG, O’Sullivan PS, Lindstaedt B. Improving graduate education to support a branching career pipeline: recommendations based on a survey of doctoral students in the basic biomedical sciences. CBE Life Sci Educ. Fall 2011;10(3):239–49. doi:10.1187/cbe.11-02-0013

13. Mathur A, Cano A, Kohl M, et al. Visualization of gender, race, citizenship and academic performance in association with career outcomes of 15-year biomedical doctoral alumni at a public research university. PLoS One. 2018;13(5):e0197473. doi:10.1371/journal.pone.0197473

14. Brandt PD, Sturzenegger Varvayanis S, Baas T, et al. A cross-institutional analysis of the effects of broadening trainee professional development on research productivity. PLoS Biol. Jul 2021;19(7):e3000956. doi:10.1371/journal.pbio.3000956

15. Gibbs KD, Jr., McGready J, Bennett JC, Griffin K. Biomedical Science Ph.D. Career Interest Patterns by Race/Ethnicity and Gender. PLoS One. 2014;9(12):e114736. doi:10.1371/journal.pone.0114736

16. Sauermann H, Roach M. Science PhD career preferences: levels, changes, and advisor encouragement. PLoS One. 2012;7(5):e36307. doi:10.1371/journal.pone.0036307

17. Wood CV, Jones RF, Remich RG, et al. The National Longitudinal Study of Young Life Scientists: Career differentiation among a diverse group of biomedical PhD students. PLoS One. 2020;15(6):e0234259. doi:10.1371/journal.pone.0234259

18. Lenzi RN, Korn SJ, Wallace M, Desmond NL, Labosky PA. The NIH "BEST" programs: Institutional programs, the program evaluation, and early data. FASEB J. Mar 2020;34(3):3570–3582. doi:10.1096/fj.201902064

19. Ullrich LE, Ogawa JR, Jones-London MD. Factors That Influence Career Choice among Different Populations of Neuroscience Trainees. eNeuro. May-Jun 2021;8(3)doi:10.1523/ENEURO.0163-21.2021

20. Kahn S, Ginther DK. The impact of postdoctoral training on early careers in biomedicine. Nat Biotechnol. Jan 10 2017;35(1):90–94. doi:10.1038/nbt.3766

21. Roach M, Sauermann H. The declining interest in an academic career. PLoS One. 2017;12(9):e0184130. doi:10.1371/journal.pone.0184130

22. Lu J, Velten B, Klaus B, Ramm M, Huber W, Coulthard-Graf R. PhD and postdoc training outcomes at EMBL: changing career paths for life scientists in Europe. bioRxiv. 2022:2022.03.01.481975. doi:10.1101/2022.03.01.481975

23. Gibbs KD, Jr., McGready J, Griffin K. Career Development among American Biomedical Postdocs. CBE Life Sci Educ. Winter 2015;14(4):ar44. doi:10.1187/cbe.15-03-0075

24. Golde CD, T. At cross purposes: What the experiences of doctoral students reveal about doctoral education. A report prepared for The Pew Charitable Trusts. Philadelphia, PA. 2001:63.

25. Lent RW, Brown SD. Social Cognitive Approach to Career Development: An Overview. The Career Development Quarterly. 1996;44(4):310–321. doi:10.1002/j.2161-0045.1996.tb00448.x

26. Lambert WM, Wells MT, Cipriano MF, Sneva JN, Morris JA, Golightly LM. Career choices of underrepresented and female postdocs in the biomedical sciences. Elife. Jan 3 2020;9doi:10.7554/eLife.48774

27. Chatterjee D, Jacob GA, Varvayanis SS, et al. Career self-efficacy disparities in underrepresented biomedical scientist trainees. PLoS One. 2023;18(3):e0280608. doi:10.1371/journal.pone.0280608

28. Bernstein R. Yes, you can attend that career event, says the U.S. government. Science Careers. 2014. doi:10.1126/science.caredit.a1400315 https://www.science.org/content/article/yes-you-can-attend-career-event-says-us-government#

29. National Institutes of Health. NOT-OD-15-008: OMB Clarifies Guidance on the Dual Role of Student and Postdoctoral Researchers. https://grants.nih.gov/grants/guide/notice-files/NOT-OD-15-008.html

30. Collins TRL, Hoff K, Starback M, Brandt PD, Holmquist CE, Layton RL. Creating and sustaining collaborative multi-institutional industry site visit programs: a toolkit. F1000Res. 2022;9:1317. doi:10.12688/f1000research.26598.2

31. Colby JM, Wheeler FC, Petrie KA, Gould KL, Schmitz JE. Institutional Training Opportunities for PhD Students in Laboratory Medicine: An Unmet Career Development Need? J Appl Lab Med. Mar 1 2020;5(2):412–416. doi:10.1093/jalm/jfz028

32. Van Wart A, O’Brien TC, Varvayanis S, et al. Applying Experiential Learning to Career Development Training for Biomedical Graduate Students and Postdocs: Perspectives on Program Development and Design. CBE Life Sci Educ. Sep 2020;19(3):es7. doi:10.1187/cbe.19-12-0270

33. Infante LL, Daniel L, Chalkley R. BEST: Implementing Career Development Activities for Biomedical Research Trainees. 1st ed. Academic Press; 2020:282.

34. Chatterjee D, Ford JK, Rojewski J, Watts SW. Exploring the Impact of Formal Internships on Biomedical Graduate and Postgraduate Careers: An Interview Study. CBE Life Sci Educ. Jun 2019;18(2):ar20. doi:10.1187/cbe.18-09-0199

35. Hartley LM, Ferrara MJ, Handelsman MM, Rutebemberwa A, Wefes I. Principles and Strategies for Effective Teaching: A Workshop for Pre- and Postdoctoral Trainees in the Biomedical Sciences. J Microbiol Biol Educ. 2019;20(3)doi:10.1128/jmbe.v20i3.1689

36. Petrie KA, Carnahan RH, Brown AM, Gould KL. Providing Experiential Business and Management Training for Biomedical Research Trainees. CBE Life Sci Educ. Fall 2017;16(3)doi:10.1187/cbe.17-05-0074

37. *National Academy of Sciences; National Academy of Engineering; Institute of Medicine*. The Postdoctoral Experience Revisited. The National Academies Press; 2014:122.

38. Yang L, Webber KL. A decade beyond the doctorate: the influence of a US postdoctoral appointment on faculty career, productivity, and salary. Higher Education. 2015;70(4):667–687.

39. Sauermann H, Roach M. SCIENTIFIC WORKFORCE. Why pursue the postdoc path? Science. May 6 2016;352(6286):663–4. doi:10.1126/science.aaf2061

40. Stephan P. How to Exploit Postdocs. BioScience. 2013;63(4):245–246. doi:10.1525/bio.2013.63.4.2

41. Konig J. Postdoctoral employment and future non-academic career prospects. PLoS One. 2022;17(12):e0278091. doi:10.1371/journal.pone.0278091

42. Langin K. U.S. labs face severe postdoc shortage. Science. 2022;376(6600):1369–1370. doi:https://doi.org/10.1126/science.add6184

43. Woolston C. Lab leaders wrestle with paucity of postdocs. Nature. Aug 30 2022;doi:10.1038/d41586-022-02781-x

44. Dunbar CEL, R.L.; Wolberg, A.S. The Perfect Storm: The Workforce Crunch and the Academic Laboratory. The Hematology. 2022;19(3) doi: https://doi.org/10.1182/hem.V19.3.2022314

45. Dzirasa K. U.S. Must Invest in Emerging Scientists. Inside HigherEd. 2023. Accessed June 9, 2023. https://www.insidehighered.com/opinion/views/2023/05/24/us-must-invest-emerging-scientists

46. Sinche M, Layton RL, Brandt PD, et al. An evidence-based evaluation of transferrable skills and job satisfaction for science PhDs. PLoS One. 2017;12(9):e0185023. doi:10.1371/journal.pone.0185023

47. Mathur A, Wood ME, Cano A. Mastery of Transferrable Skills by Doctoral Scholars: Visualization using Digital Micro-credentialing. Change. 2018;50(5):38–45. doi:10.1080/00091383.2018.1510261

48. Braun V, Clarke V. Using thematic analysis in psychology. Qualitative Research in Psychology. 2006;3:77–101. doi:10.1191/1478088706qp063oa

49. Strengthening the Biomedical Research Workforce. National Institutes of Health. Office of Strategic Coordination - The Common Fund. Accessed May 9, 2023, https://commonfund.nih.gov/workforce/programresources

50. R Core Team. R: a language and environment for statistical computing. https://www.R-project.org/

51. Zeileis AH, T. Diagnostic Checking in Regression Relationships. R News. 2002;2(3):7–10.

52. Wickham H. ggplot2: Elegant Graphics for Data Analysis. 2nd ed. Use R! Springer-Verlag; 2016.

53. Walters SJ, Campbell MJ, Machin D. Medical Statistics: A Textbook for the Health Sciences. 5th ed. Hoboken: Wiley-Blackwell; 2021:448.

