## Supplemental appendix for "From goal to outcome: analyzing the progression of biomedical sciences PhD careers in a longitudinal study using an expanded taxonomy"

**S1 Appendix:** 3-Tiered Taxonomy and Flag System for Classifying Biomedical Sciences PhD Alumni (v. 2019)

#### Overview of the record classification process

The Office of Biomedical Research Education and Training (BRET) at Vanderbilt University School of Medicine classifies each alumni employment record using a 3-tiered taxonomy and series of "flags" intended to capture additional characteristics about each job. The Sector and Career Type tiers were modeled on how STEM PhD careers are categorized in the *NSF Survey of Doctoral Recipients*. Our initial list of Job Functions was developed by grouping similar careers together, then subsequently modified for greater consistency with the classifications used by the NIH BEST consortium. The flags are a series of fields with limited answer options that record additional characteristics of each job. These flags are useful for classifying complex employment situations and denoting important career milestones.

| | 3-tiered taxonomy | | | Flags | | | | | | | |
|---|---|---|---|---|---|---|---|---|---|---|---|
| | **Sector** | **Career Type** | **Job Function** | **Employment status** | **Primary/ Secondary** | **Science - related** | **1st desti-nation after PhD** | **1st non-training position** | **Entre-preneur-ship** | **Execu-tive** | **Faculty appoint-ment** |
| **Answer options** | *See S1 Appendix pages 3-9 for answer options* | | | Employed<br>Not Employed<br>Unknown | Primary<br>Secondary | Yes<br>No<br>Unknown | Yes<br>No | Yes<br>No<br>Unknown | Yes<br>No<br>Unknown | Yes<br>No<br>Unknown | Yes<br>No<br>Unknown |

Our 3-tiered taxonomy scheme is similar, but not identical to, the taxonomy published by Mathur et al. (2018) and refined by Stayart et al. (2020). Like Mathur et al. and Stayart et al., our taxonomy categorizes each job by Sector, Career Type, and Job Function. However, two notable differences in our taxonomy are that we do not define careers as "science-related" or "not related to science" in the "Career Type" tier, and we do not use tenure or part-time status to define different types of faculty careers in the "Job Function" tier. Instead, our taxonomy uses a series of flags to indicate whether *every* job is science-related or carries a faculty appointment. Rationale for the flags and faculty categorization system is described below.

#### Rationale for each flag

- Employment status: "One-touch" option to remove alumni who are unemployed, thus facilitating analysis of the distribution of *employed* alumni among sectors, career types, or job functions.
- Primary/secondary: Allows us to capture employment information for alumni who simultaneously hold multiple jobs. For example, a professor with a start-up would have two employment rows, one for professor (primary) and a second for start-up founder (secondary).
- Science-related: Some taxonomies categorize "Not science related" jobs as a "Career Type" in the 3-tiered taxonomy. However, this creates classification conflicts because "Not science related" is not mutually exclusive with other Career Types. For example, someone who teaches IP law at a law school is doing a job that is "Primarily teaching" AND (arguably) "Not science related." To avoid these classification conflicts, we classify every position in the 3-tiered taxonomy and use a flag to mark jobs "Not science related."
- First destination after PhD: About 30% of our alumni do brief (1-6 months) postdoctoral fellowships in their PhD lab, often to finish a paper or time their move. Sometimes these alumni subsequently start another postdoc and sometimes they start a job. The "first destination after PhD" field allows us to designate their *subsequent* position after a short postdoc in their PhD lab as their "First destination after PhD," providing for more accurate assessment of the number of alumni who do postdocs that are distinct from their PhD research.
- First non-training position: Useful for comparing first non-training positions across alumni, regardless of when the position was obtained.
- Entrepreneurship: Distinct from the "Entrepreneurship" Job Function, which is reserved for alumni who start companies based on scientific discoveries or healthcare technologies, this flag denotes individuals who start companies of any type, including services like consulting.
- Executive: Identifies alumni in high-level positions of leadership within their organizations.
- Faculty appointment: see next page

**Faculty Classification System**

Classifying faculty positions is complex. Titles like, “faculty” or “professor” do not convey the type of work someone does (e.g. research vs. teaching vs. clinical work) nor do they convey characteristics like tenure-status or temporary appointment. Some taxonomies deal with this complexity by having multiple faculty Job Functions that are various combinations of type of work and tenure status. However, we found this approach challenging because there are four permutations of the most common categories (research vs. teaching + nontenure vs. tenured/tenure track) and we often encountered alumni who didn't fit neatly into any of these four options, e.g. alumni whose faculty role was primarily administrative or clinical, and alumni whose faculty appointment was clearly part-time or temporary.

To deal with this complexity, our taxonomy categorizes faculty in the Job Function tier according to their primary job function, most often, “principal investigator or group leader” or “research faculty or staff” or “teaching faculty or staff” or “administration” or “healthcare.” Records of individuals who have a faculty appointment are then assigned a faculty flag, “faculty appointment=yes,” and classified further. We record three elements of the job title (prefix, rank, and title) and assign a presumed tenure status to each record. Recording the prefix, rank, and faculty title is straightforward and allows us to follow the progression of alumni faculty careers over time. Furthermore, certain faculty titles or prefixes are presumed indicators of nontenure status, e.g. “adjunct” or “visiting” or “research.”

| Field | Answer Options | Rules |
|---|---|---|
| Faculty Prefix | Adjunct, Clinical, Visiting, Research, Other | If there is no prefix in the individual’s job title >>> leave field blank.<br>Use “Other” for all other prefixes that are not defined here.<br>If prefix = “adjunct” or “visiting” >>> presumed tenure status = “tenure not applicable.”<br>If prefix = “research” or “clinical” >>> presumed tenure status = “nontenure track.” |
| Faculty Rank | Assistant, Associate, Other | If there is no rank in the individual’s job title >>> leave field blank. |
| Faculty Title | Instructor, Lecturer, Professor, Other | If faculty title = “instructor” or “lecturer” >>> presumed tenure status = “nontenure track.” |
| Presumed Tenure Status | Foreign Institution<br>Nontenure Track<br>Tenured or Tenure Track<br>Tenure Not Applicable<br>Tenure Status Unclear | Faculty at non-US universities are always designated, “Foreign institution.”<br>If none of the above prefix/rank/title rules apply, then “presumed tenure status” is assigned based on publicly available information. Evidence of tenure-track or tenured status includes:<br>• Announcements of tenure or promotion on university websites or social media<br>• Departmental listings of faculty, often with a link to faculty CVs or research programs<br>• Website describing the faculty member’s research program and team<br>• Named professorships or leadership positions within the department<br>• Presence of large research or educational grants from major funders like the NIH or NSF<br>• Multiple last-author publications (biomedical sciences)<br>Evidence of nontenure track status is a lack of the above indicators, as well as being listed exclusively as part of a research core facility or another faculty member’s research group.<br>If our confidence in assigning tenure status is low after reviewing publicly available information, the record is assigned “tenure status unclear.” This most often occurs for brand new institutions or faculty appointments, and this field is updated if tenure status becomes clear at a later timepoint. |

### 3-Tiered Taxonomy

| Tier 1: Sector | Answer options | Definition | Coding clarifications |
|---|---|---|---|
| | Academia | Any academic institution where education or research training occurs including colleges, universities, K-12 schools, and some medical centers with "university" in the name. | Includes healthcare clinics, hospitals, and medical centers that have the word "university" in the name of the organization. If "university" is not in the name of the healthcare provider or hospital, the record should be classified in a different sector (usually nonprofit). Except for K-12 schools, foreign universities, and newly established universities, all records that are categorized as "sector = Academia" should have a Carnegie ID |
| | Government | Organizations operated by federal, state, local or foreign governments. | Includes VA hospitals, national laboratories, and military laboratories |
| | For-Profit | Organizations or entities that operate to make a profit, including some industry research. | Includes self-employed individuals, unless they operate not-for-profit entities |
| | Nonprofit | Non-governmental organizations that do not operate to make a profit. | Includes many hospitals, healthcare centers, and independent research institutes (see Association of Independent Research Institutes for a list). If the organization has the word "university" in the name, it is coded as Academia |
| | Not employed | For individuals who are not currently employed or unemployed | Includes individuals who are known to be out of the workforce as full-time caretakers or parents |
| | Deceased/Retired | For individuals who are deceased or retired | |
| | Unknown | For individuals for whom no employment information is found or known. | |

#### Tier 2: Career Type

| Answer options | Definition | Coding clarifications |
|---|---|---|
| Primarily research | The primary, although not necessarily the only, focus is the conduct or oversight of scientific research. | Most faculty at US research-intensive institutions with very high research activity, or medical schools, as identified through the 2018 Carnegie classifications 15 or 25, will be included here. Individuals whose primary role is to conduct research will usually oversee or be a member of a research team. |
| Primarily teaching | The primary, although not necessarily the only, focus is education and teaching. | Includes most academic faculty at all other Carnegie institutions that are not mentioned above. |
| Primarily administrative or managerial or operational | The primary, although not necessarily the only, focus is administrative functions, managing people or programs, or overseeing or executing operational functions. | Does NOT include individuals who oversee research activities. Healthcare providers, except medical residents or individuals in training, should be included here; residents, interns, and others in training should be classified in "further education or training." |
| Further education or training | Temporary education or training position *other than* research postdoctoral fellows. | Includes individuals who are completing medical residency, interns, and those pursuing an additional degree. Postdoctoral fellows who are conducting research should be classified as "primarily research." Other types of "fellows" (clinical, policy, data science, regulatory) should be classified here. |
| Not employed | For individuals who are not currently employed or unemployed | Includes individuals who are known to be out of the workforce as full-time caretakers or parents |
| Deceased/Retired | For individuals who are deceased or retired | |
| Unknown | For individuals for whom no employment information is found or known. | |

#### Tier 3: Job Function

| Answer options | Definition | Coding clarifications | Example duties/titles |
|---|---|---|---|
| Administration | Role that involves the administration of science, research, or education-related programs or products in academia or nonprofit sectors | Many academic and nonprofit administration jobs go here, including executive leadership. Individuals in administrative roles in government should be coded as job function="science policy or government affairs." Generally, individuals in administrative roles in for-profit companies should be coded elsewhere according to the division of the company in which they work. | Faculty affairs, academic program administrators, human resources, academic admissions, career development offices, grant and contracts management, research development, PhD-level program development, project manager, dean, provost, chancellor, department chair, center director, assistant dean, associate dean |
| Business development, consulting, and strategic alliances | Role that involves the development, execution, management, or analysis of a business. Role may include relationship management, refinement of operational efficiency, or fee-based advisory services. | Includes subject matter experts, technical consultants, competitive intelligence, and third-party market analysis. Does not include chief scientific officer (CSO), which is classified as "Principal investigator or group leader" with an Executive flag. Does not include academic deans, who should be classified in Administration with Executive flag. | Management consultant, business development professional, market researcher, investment analyst, venture capitalist |
| Clinical research management or clinical development | Role that is responsible for the administrative management of clinical research trials or patients or data, planning or execution of clinical trials, or clinical development of compounds | These positions may involve coordination with regulatory and scientific groups. Some quality assurance (QA) roles may fit well here. Other QA roles may fit better under regulatory affairs or manufacturing, depending on the context. | Clinical research project manager, clinical research trials manager or coordinator, clinical data manager, clinical data analyst, director of clinical development |
| Clinical services | Role that involves the execution or interpretation of diagnostic or forensic tests or administration of clinical services, not including healthcare providers. | Does not include healthcare providers or clinical research/trials-related work. | Genetic counselor, testing specialist, clinical laboratory directors or staff, forensic scientist, neurophysiological monitoring |

#### Tier 3: Job Function (cont.)

| Answer options | Definition | Coding clarifications | Example duties/titles |
|---|---|---|---|
| Completing further education | Pursuing additional education that results in graduation with conferment of a degree or certificate. | Only degree or certificate-granting programs should go here. Research postdoctoral fellowships should be classified as "Postdoctoral." Other types of training programs (internships, residencies, non-traditional postdocs in policy, regulatory affairs, data science, etc.) should be classified in the job function that best captures the type of work. For example, a resident should be classified as, "healthcare provider" and an AAAS policy fellow should be classified as "science policy or government affairs." | Law student, medical student, veterinary student, nursing student, MBA student |
| Data science, analytics, and software engineering | Role that may combine programming, analytics, advanced statistics, data communication, and/or software development. | Accompanying career type (research OR administrative/operational) determined by type of work they do; data scientists should be classified as "career type=primarily research" and programmers and software engineers should be classified as "career type=primarily administrative, managerial, or operational." | Programmer, data scientist, software architect or engineer, genomic data analyst |
| Entrepreneurship (scientific or healthcare) | Founder or co-founder of a company that develops, manages, and provides/obtains capital to initiate a business or enterprise based on a scientific, research, or healthcare discovery or product. | Does not include C-suite of well-established companies or C-suite employees who were not involved in establishing the original business; these individuals should be classified in other categories, likely as "business development". Does not include staff at start-ups, or individuals who are sole proprietors providing an administrative service (e.g., medical writers, consultants). | Founder, co-founder |

### Tier 3: Job Function (cont.)

| Answer options | Definition | Coding clarifications | Example duties/titles |
|---|---|---|---|
| Healthcare provider | Role where the primary responsibility is providing healthcare | | Doctor, nurse, medical resident, clinical fellow, veterinarian, dentist |
| Intellectual property and law | Role that involves the curation, management, implementation or protection of intelligence and creation, including trademarks, copyrights, patents, or trade secrets. | Includes technology transfer specialists at universities, nonprofits, and government agencies. Includes employees of the US Patent and Trademark Office. | Patent agent, patent attorney, patent scientist, patent reviewer, technology transfer specialist. |
| Manufacturing and process development | Role that involves the oversight, design, or optimization of manufacturing processes | Some quality assurance roles may fit well here. Others may fit better under regulatory affairs or clinical development, depending on the context. | Process engineer, quality assurance engineer |
| Medical affairs | Role that involves providing scientific and clinical support for commercial products. | Does not include medical writers, who should be classified under "scientific or medical writing or communication" | Medical science liaison, medical affairs officer or manager, medical director, medical information officer |
| Postdoctoral | Temporary mentored research training position following completion of doctoral degree. | Use only for postdocs that appropriately coded as "career type" = Primarily Research. | |
| Principal investigator or group leader | Role in an academic, industry, government, or nonprofit setting where the primary responsibility is leading a basic, translational, or preclinical research team or program. | These individuals are responsible for funding and/or managing a research program or group(s). Role is directly responsible for determining the direction of research programs. Titles such as Principal Scientist and Director need to be evaluated on a case-by-case basis and, if similar to academic core facility directors, these titles should be classified as "research staff or technical director." Generally, it does NOT include Scientist I/II/III or Senior Scientist titles. | Principal investigator (PI), tenured and tenure-track "professor" titles at universities, research institutes, VA hospitals, and other government research institutions, chief scientific officer (CSO), director of research & development, group leader, vice president of research. |

### Tier 3: Job Function (cont.)

| Answer options | Definition | Coding clarifications | Example duties/titles |
|---|---|---|---|
| Regulatory affairs | Role that involves controlling or evaluating the safety and efficacy of products in areas including pharmaceuticals, medicines, and devices | Any science/medical writing that is regulatory in nature should be classified here; Some quality assurance roles may fit well here. Others may fit better under manufacturing or clinical development, depending on the context. | Institutional regulatory affairs professional, quality control specialist, compliance officer, senior safety specialist |
| Research staff or technical director | Role that directly involves performing or managing research | This should be used unless the individual is responsible for managing an independent research group. Nontenure track research faculty are included here. Most entry-level scientist roles in industry are included here. | Senior scientist, staff scientist, scientist I/II/III, lab/core managers, directors of research facilities, public health analyst, epidemiologists, research assistant professor, research instructor, research associate |
| Sales and marketing | Role that is related to the sales or marketing of a science-related product or service | This does not include medical science liaisons, who should be coded under medical affairs | technical sales representative, marketing specialist |
| Science education and outreach | Role that involves K-12 teaching or public outreach at a primary/secondary school, science museum, scientific society, or similar | Includes primary and secondary education teachers (K-12 teachers) | K-12 educator, high school teacher, museum curriculum development, outreach program administrator |
| Science policy and government affairs | Role that involves policy or program development, operations, or review, including analysis, advisory, or advocacy | All non-research positions in government are coded here. | Program officer, public affairs or government affairs staff at scientific societies, foundations, government entities, or think tanks; Science Policy Analyst, Public Health Analyst, AAAS Science & Technology Policy fellow |

#### Tier 3: Job Function (cont.)

| Answer options | Definition | Coding clarifications | Example duties/titles |
|---|---|---|---|
| Science or medical writing or communication | Role that involves the communication of science-related topics | Does not include regulatory writing, which should be classified under "regulatory affairs" | Science, medical, or technical writer, journalist, science editor, science publisher |
| Teaching faculty or staff | Post-secondary teaching position where teaching is majority of the individual's professional activity. | This may include faculty at teaching-intensive institutions who maintain research programs primarily for the purpose of educating undergraduates; the majority of the time/effort and the salary of individuals in this category typically comes from the institution and not research grants. K-12 teachers are NOT coded here; they should go under STEM education and outreach. | |
| Technical support and product development | Role that requires specialized technical knowledge of a science-related product | | Technical support specialist, field application specialist, product development scientist or engineer |
| Other | Some other job function that is not defined elsewhere | These positions may be science-related or non-science related | Service sector jobs; ownership of non-science related companies |
| Not employed | For individuals who are not currently employed or unemployed | | Includes individuals who are known to be out of the workforce as full-time caretakers or parents |
| Deceased/retired | For individuals who are deceased or retired | | |
| Unknown | For individuals for whom no employment information is found or known. | | |
