## Supplemental table 1 for "From goal to outcome: analyzing the progression of biomedical sciences PhD careers in a longitudinal study using an expanded taxonomy"

**S1 Table: Representative job titles in each career area**

| | |
|---|---|
| **Academic Research (Ac Research)**<br>*These positions occur in the academic sector.* | Postdoctoral Fellow, Assistant Professor or higher ranks, Research Assistant Professor or higher ranks, Research Instructor, Staff Scientist, Senior Research Associate, Research Specialist, Project Manager, Imaging Facility Manager |
| **For-profit Research (Fp Research)**<br>*These positions occur in the for-profit sector.* | Postdoctoral Fellow, Senior Scientist, Principal Scientist, Chief Scientific Officer (CSO), VP Biology, Senior Director, R&D Manager, Principal Statistician, Director Small Molecule Screening, Project Manager, Associate Director |
| **Government or Nonprofit Research (G/Np Research)**<br>*These positions occur in the government or nonprofit sectors.* | Postdoctoral Fellow, Research Associate, Staff Scientist, Assistant Professor, Principal Investigator, Supervisory Biologist, Research Health Scientist, Laboratory Manager, Interdisciplinary Scientist, Biostatistician, Associate Member, Analytical Sciences Group Leader |
| **Teaching**<br>*These positions occur in the academic sector in higher education institutions or K-12 schools.* | Assistant Professor or higher ranks, Instructor, Senior Lecturer, Adjunct Faculty, Visiting Assistant Professor, High School Teacher, Science Teacher |
| **Administrative, Managerial, or Operational (AMO)**<br>*These positions can occur in any sector – academia, private industry, government, or nonprofit organizations.* | Medical Writer, Medical Science Liaison, Physician, Patent Agent, Product Manager, Scientific Advisor, Manager of Business Development, Vice President of Business Intelligence, Technology Commercialization Associate, Staff Software Engineer, Regulatory Affairs Specialist, Clinical Research Associate, Account Manager, Life Sciences Consultant, Director of Clinical Research Operations, Scientific Review Officer, Program Manager, Director, Project Manager, Science Officer, Patent Examiner, Intelligence Analyst, Health Scientist Administrator, Associate Dean for Academic Affairs |
| **Further Education or Training**<br>*These positions are typically in the academic sector, though some are in government or nonprofit organizations.* | Medical Student, Law Student, MBA Student, Veterinary Student, Nursing Student, MS Student, MPH Student, Dental Student, Resident, AAAS Science & Technology Policy Fellow, Presidential Management Fellow, FDA Commissioner's Fellow |
