## Supplemental table 2 for "From goal to outcome: analyzing the progression of biomedical sciences PhD careers in a longitudinal study using an expanded taxonomy"

**S2 Table: Career area at Y1, Y3, and Y5 compared to Y10**

| | | Career area at Y10 | | | | | | Total |
|---|---|---|---|---|---|---|---|---|
| | | Ac Research | Fp Research | G/Np Research | Teaching | AMO | Further Education | |
| Career area at Y1 | Ac Research | 172 | 61 | 33 | 43 | 108 | 1 | 418 |
| | Fp Research | 1 | 15 | 1 | 1 | 8 | | 26 |
| | G/Np Research | 29 | 16 | 10 | 3 | 26 | 1 | 85 |
| | Teaching | 2 | - | - | 20 | 3 | - | 25 |
| | AMO | - | 1 | - | - | 60 | - | 61 |
| | Further Education | 3 | - | - | - | 30 | 6 | 39 |
| Career area at Y3 | Ac Research | 168 | 43 | 24 | 29 | 61 | 1 | 326 |
| | Fp Research | 4 | 32 | 2 | 2 | 19 | - | 59 |
| | G/Np Research | 27 | 17 | 18 | 2 | 23 | 1 | 88 |
| | Teaching | 3 | - | - | 33 | 4 | - | 40 |
| | AMO | 2 | 1 | - | - | 104 | - | 107 |
| | Further Education | 3 | - | | 1 | 24 | 6 | 34 |
| Career area at Y5 | Ac Research | 175 | 29 | 10 | 16 | 32 | - | 262 |
| | Fp Research | 3 | 46 | 3 | - | 21 | - | 73 |
| | G/Np Research | 19 | 15 | 29 | 1 | 12 | 1 | 77 |
| | Teaching | 4 | - | - | 49 | 2 | - | 55 |
| | AMO | 3 | 3 | 2 | 1 | 155 | 1 | 165 |
| | Further Education | 3 | - | - | - | 13 | 6 | 22 |
